# Sulfated host glycan recognition by carbohydrate sulfatases of the human gut microbiota

**DOI:** 10.1101/2021.07.23.453482

**Authors:** Ana Luis, Arnaud Baslé, Dominic P Byrne, Gareth SA Wright, James London, Jin Chunsheng, Niclas Karlsson, Gunnar Hansson, Patrick Eyers, Mirjam Czjzek, Tristan Barbeyron, Edwin A Yates, Eric C. Martens, Alan Cartmell

## Abstract

The vast microbial community that resides in the human colon, termed the human gut microbiota, performs important roles in maintaining host health. Sulfated host glycans comprise both a major nutrient source and important colonisation factors for this community. Carbohydrate sulfatases remove sulfate groups from glycans and are essential in many bacteria for the utilisation of sulfated host glycans. Additionally, carbohydrate sulfatases are also implicated in numerous host diseases, but remain some of the most understudied carbohydrate active enzymes to date, especially at the structural and molecular level. In this work, we analyse 7 carbohydrate sulfatases, spanning 4 subfamilies, from the human gut symbiont *Bacteroides thetaiotaomicron*, a major utiliser of sulfated host glycans, correlating structural and functional data with phylogenetic and environmental analyses. Together, these data begin to fill the knowledge gaps in how carbohydrate sulfatases orchestrate sulfated glycan metabolism within their environment.

## Introduction

The human gut microbiota (HGM) is a vast microbial community^1^ that makes important contributions to its host’s physiology by contributing to an array of biological processes, including protection from pathogens^2^, regulating the immune system^3^, and providing up to 10 % of the hosts energy needs through complex carbohydrate fermentation^4^. The Bacteroidetes constitute a major phylum within this community. These bacteria thrive within this competitive environment by metabolising complex glycans derived from both the diet, namely plant cell wall polysaccharides and other dietary fibers^5–7^, and the host, which continuously secretes glycan-rich substances such as mucins found in the protective mucus layers of the body^8–10^. Some *Bacteroides* dedicate as much as 20 % of their genome to glycan metabolism^11, 12^ and members of this phylum arrange their CArbohydrate Active enZymes (CAZymes) into polysaccharide utilisation loci (PUL)^13^, which are sets of genes that are genetically co-localised and co-regulated in response to a particular glycan.

The ability to metabolise sulfated host glycans such as glycosaminoglycans (GAGs), which include heparan sulfate (HS)^8^ and chondroitin sulfate (CS)^9^, and colonic mucin *O*-glycans^10^ (**Figure 1**) has been shown for several *Bacteroides* species. These complex carbohydrates are also important colonisation factors^14, 15^, and GAGs have also been shown to be high priority nutrients^16, 17^. Thus, the effects of host glycans on the microbiota composition could be profound. Carbohydrate sulfatases are enzymes that catalyse the de-sulfation of host glycans and are essential for their utilisation^8^ by bacteroides species of the HGM. Additionally, carbohydrate sulfatases produced by gut microbes have been implicated in inflammatory bowel disease (IBD) in humans^18, 19^, and directly linked to promoting colitis in a susceptible mouse model^20, 21^. A recent study has also demonstrated that carbohydrate sulfatase activity is a keystone step in colonic mucin metabolism and their loss results in an inability of at least one *Bacteroides* species to utilise colonic mucin and effectively colonise the mouse gut^10^. The model organism *Bacteroides thetaiotaomicron* (*B. theta*) possesses at least 28 sulfatases spread amongst several PULs of both known, and unknown, function in glycan degradation^8–10^. Despite the important role of carbohydrate sulfatases in glycan degradation, particularly host glycans, there is a significant knowledge gap regarding the structural basis behind the substrate recognition by these enzymes.

**Figure 1.**
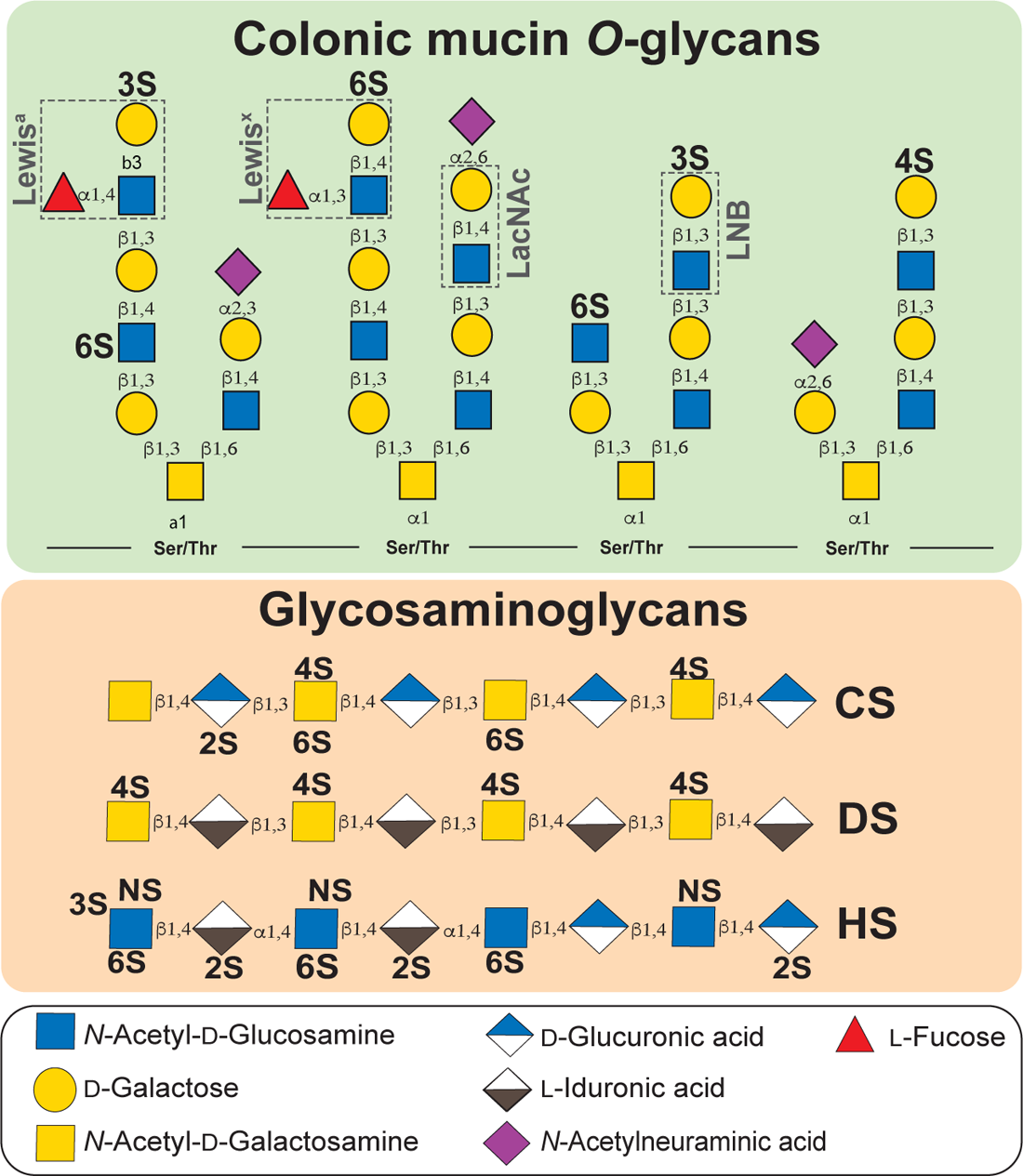
Examples of sulfated host carbohydrates found in the colon. Schematic representation of sulfated host glycans found in mucin O-glycans (green box) and glycosaminoglycans which are integral to the extracellular matrix and glycocalyx (peach box), representing a constant, host derived, nutrient source for the colonic microbiota. Sugars are shown according to the Symbol Nomenclature for Glycan system^47^.

Sulfatases are catalogued in the SulfAtlas database (http://abims.sb-roscoff.fr/sulfatlas/)^22^. These enzymes are currently divided into four families, S1-S4, based on sequence similarity, fold, and catalytic mechanism. The S1 family is the largest family comprising ∼90 % of all sulfatase sequences, is found throughout all domains of life, and is currently the only family that has been significantly implicated in carbohydrate metabolism. There are currently 72 S1 subfamilies (denoted as S1_X) comprising a total of 36,815 individual sulfatases yet less than 1 % have had their activities analysed and only ∼10 unique carbohydrate sulfatase structures exist. This makes S1 carbohydrate sulfatases some of the most poorly characterised CAZymes to date.

The S1 family is part of the alkaline-phosphatase-like superfamily. All S1 sulfatases require either a Cys or Ser residue, within the core consensus sequence **C/S**-X-P/A-S/X-R, to be co-transitionally transformed into formylglycine (FGly) to be catalytically active^23^. Calcium is an essential cofactor for all S1 sulfatases, whilst the catalytic acid is likely an invariant His or Lys that coordinates with the scissile sulfoester linkage. The subsite nomenclature for carbohydrate sulfatases is such that the invariant sulfate binding site is denoted the S site. The S site sulfate is appended to the 0 subsite sugar. Subsites then increase in number (i.e. +1, +2, +3) as the sugar moves toward the reducing end (free *O*1) and decreases in number as the sugar chain moves towards the non-reducing end (i.e. −1, −2, −3)^24^.

Here, we describe the structures of 7 *Bacteroides* S1 carbohydrate sulfatases, 6 in complex with ligand (**Figure S1**), along with detailed biochemical, mutagenic, and bioinformatic analyses. The sulfatases span 4 different subfamilies, with three of the structures representing the first structural description of S1_16 and S1_46 subfamilies, whilst the additional structures of S1_11 and S1_15 subfamilies reveal how non-conserved areas of the proteins are adapted to increase specificity for the particular glycan targeted by the PUL in which they reside. The data reveal the exquisite specificity that S1 carbohydrate sulfatases utilise to recognise their cognate sulfated glycan within the human colon and lay the foundations for developing strategies to manipulate this specificity to improve microbiome health, and thus that of the host.

## Results

### Conserved features of S1 formylglycine sulfatases

Consistent with the previously characterized structures, all the S1 sulfatases investigated here adopt an α/β/α fold for the core N-terminal domain which is abutted by a smaller, C-terminal, ‘sub domain’ (**Figure 2a**). The sulfate binding site (S site) is invariant across the S1 family. The FGly residue sits at the base of the pocket in the consensus sequence **C/S**-X-P/A-S/X-R. The varying ratios of Cys/Ser at the critical FG position are 83/17 for S1_11, 82/18 for S1_15, 85/15 for S1_16 and 84/15 for S1_46 (**Figure 2b**). Pro is conserved in 95 % of the sequences except in S1_46 where Pro exists 90 % of the time with Ala being the other 10 % (**Figure 2b**). Although *B. theta* sulfatases exclusively possess Ser, the favouring of Cys over Ser reflects that only organisms inhabiting anaerobic environments are currently known to convert Ser to FGly^23^. Individual enzymes are subsequently referred by their gene/locus tag number with the corresponding activity in superscript (*e.g*., BT1918^3S-GlcNAc^). All structures solved were Ser variants and, with exception of BT1918^3S-GlcNAc^, had occupation of calcium in the metal binding site. In the S1_11, S1_15, and S1_16 subfamilies three Asp residues, a Gln/Asn and the FGly coordinate the calcium with an octahedral coordination completed by the incoming sulfate substrate (**Figure 2c**). In BT1918^3S-GlcNAc^ one of the conserved Asp is replaced by Q292 and Gln/Asn by H318 (**Figure 2c**). Q292 is positioned further away than the conserved Asp and may change the coordination to trigonal bi-pyramid and a weaker calcium binding capacity. Additionally, the absence of the FGly residue will also make calcium binding weaker in the BT1918^3S-GlcNAc^ structure. This was previously observed in BT1596^2S-Δ4,5UA^, an S1_9 sulfatase, where both Asp and Gln/Asn residues are replaced by His^8^ (**Figure 2c**)

**Figure 2.**
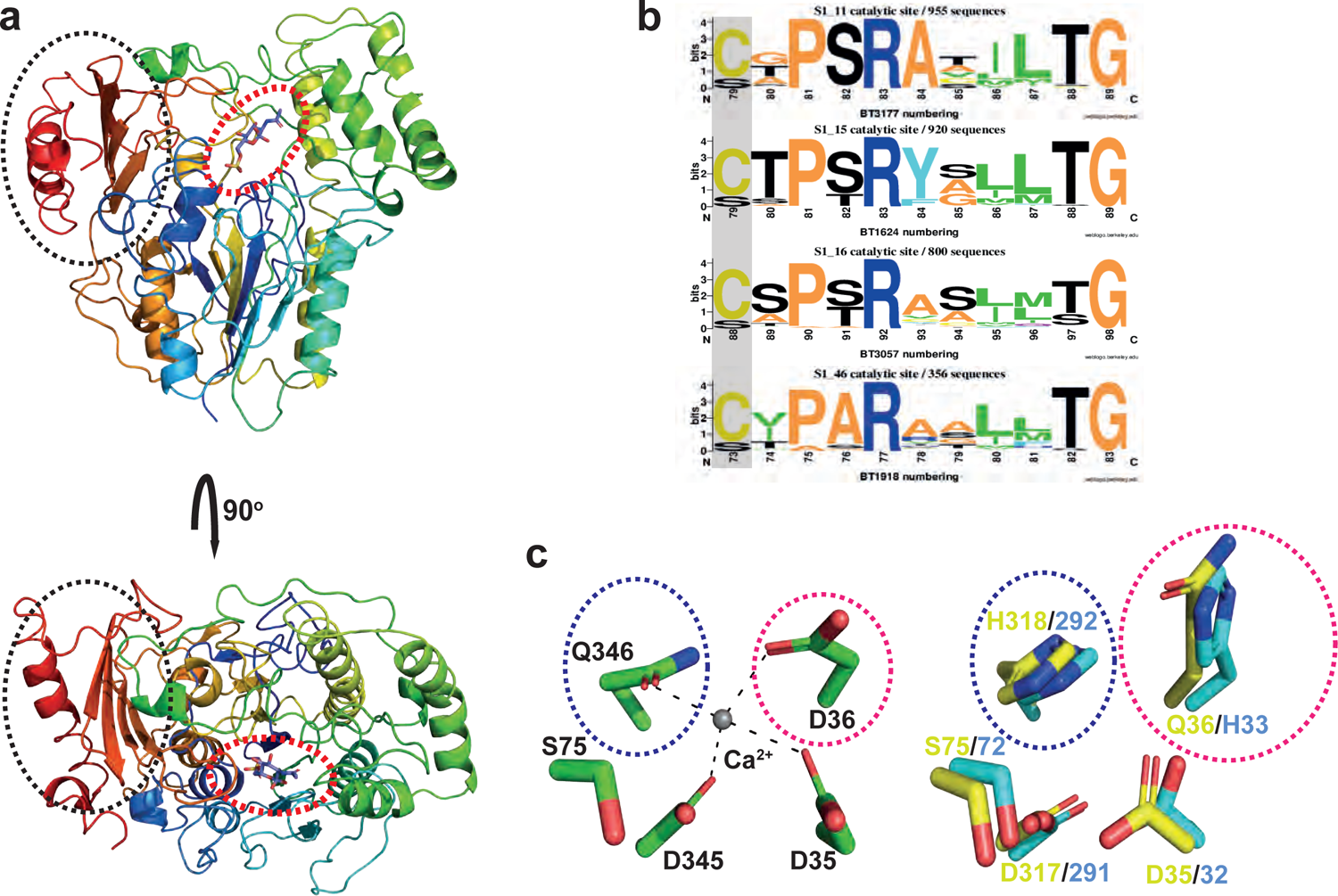
Conserved features of S1 carbohydrate sulfatases. a, Example of the overall fold common to all S1 sulfatases, Cartoon representation of BT31776S-GlcNAc colour ramped from blue (***a****/b/****a*** N-terminal domain) to red (***b***-sheet C-terminal domain). The black dashed circle indicates the C-terminal sub-domain and the red dashed circle the active site location on the core ***a/b/a*** domain. **b,** Conservation of the consensus active site sequence for S1_11, S1_15, S1_16 and S1_46. The position of formylglycine installation is highlighted with a grey box **c,** Examples of the calcium binding site: left side (green), representation of BT31776S GlcNAc calcium site as an example of the typical site observed in most sulfatase structures described to date; right side, display of alternate sites observed in BT19183S-GlcNAc (yellow) and BT15962S-*D*4, 5UA (cyan). The variable areas are highlighted in blue and red dashed circles.

### S1_46 sulfatases target rare linkages found in host glycosaminoglycans

BT1918^3S-GlcNAc^ is a dimer in solution **(Figure S2a)** and the only sulfatase identified to date that cleaves sulfate from the *O*3 position of 3S,6S-D-*N*-acetylglucosamine (3S,6S-GlcNAc)^10^, a known component of the host glycans heparin (Hep) and heparan sulfate (HS) (**Figure 1**). The *O*6 sulfate does not appear to contribute to substrate specificity, and mutation of W173 to Ala, the only residue located near the O6 sulfate, causes no significant loss in activity (**Figure 3a, 3b, S2b** and **Table S1**). Out of the five residues in the carbohydrate binding region three interact with the *N*-acetyl group (Y94, R327, and Y408) (**Figure 3a**). Mutation of Y94, Y408, and R327 individually to Ala causes a ∼40, ∼20, and ∼10-fold loss in activity, respectively (**Figure 3b, S2b, S2c,** and **Table S1**). This suggests that the interactions with *N*-acetyl group are a major specificity determinant in S1_46 sulfatases. A finding that is consistent with the lack of activity of BT1918^3S-GlcNAc^ on 3S- and 3S,6S-glucosamine^10^ (**Figure S2d**). The sulfatase activity was not affected by the mutation of Y94 and Y408 to Phe indicating their action is through the steric/stacking contributions of the phenol ring. N174 binds to the endocyclic ring oxygen via Nδ2 and R143 coordinates with O4 via Nη1, both helping to orientate the *O*3 of the sugar (**Figure 3a**). The mutation of N174 and R143 to Ala causes a ∼30 and ∼40 fold reduction in catalytic activity, respectively (**Figure 3b and Table S1**), indicating that these residues have a significant role in the substrate recognition. Phylogenetic analyses show that N174 and R143 are conserved in 87 % and 76 %, respectively, of the aligned S1_46 sequences. Additionally, Y94 and R327 are conserved in ∼65 % sequences. Y94 is present in the motif GRVGYGDE and is replaced by Asn and Phe 18 % and 10 % of the time, respectively (**Figure 3c**). Mutational data indicate that the Phe substitution does not affect activity. It is of note that Y94, N174, R327, and Y408 are invariant within Bacteroidetes inhabiting the human gastrointestinal tract (**Table S2**). In fact, the majority of sequences from S1_46 are derived from human sources and include several human pathogens, including *Clostridium difficile P28*, a sequence which contains residues equivalent to Y94, N174, and R327 but not Y408 (**Table S2**). Interestingly, Y408 is only found in 26% of S1_46 sequences, within a HATCY motif, but is present in all Gram-negative bacteria within this subfamily indicating some phylum level differences.

**Figure 3.**
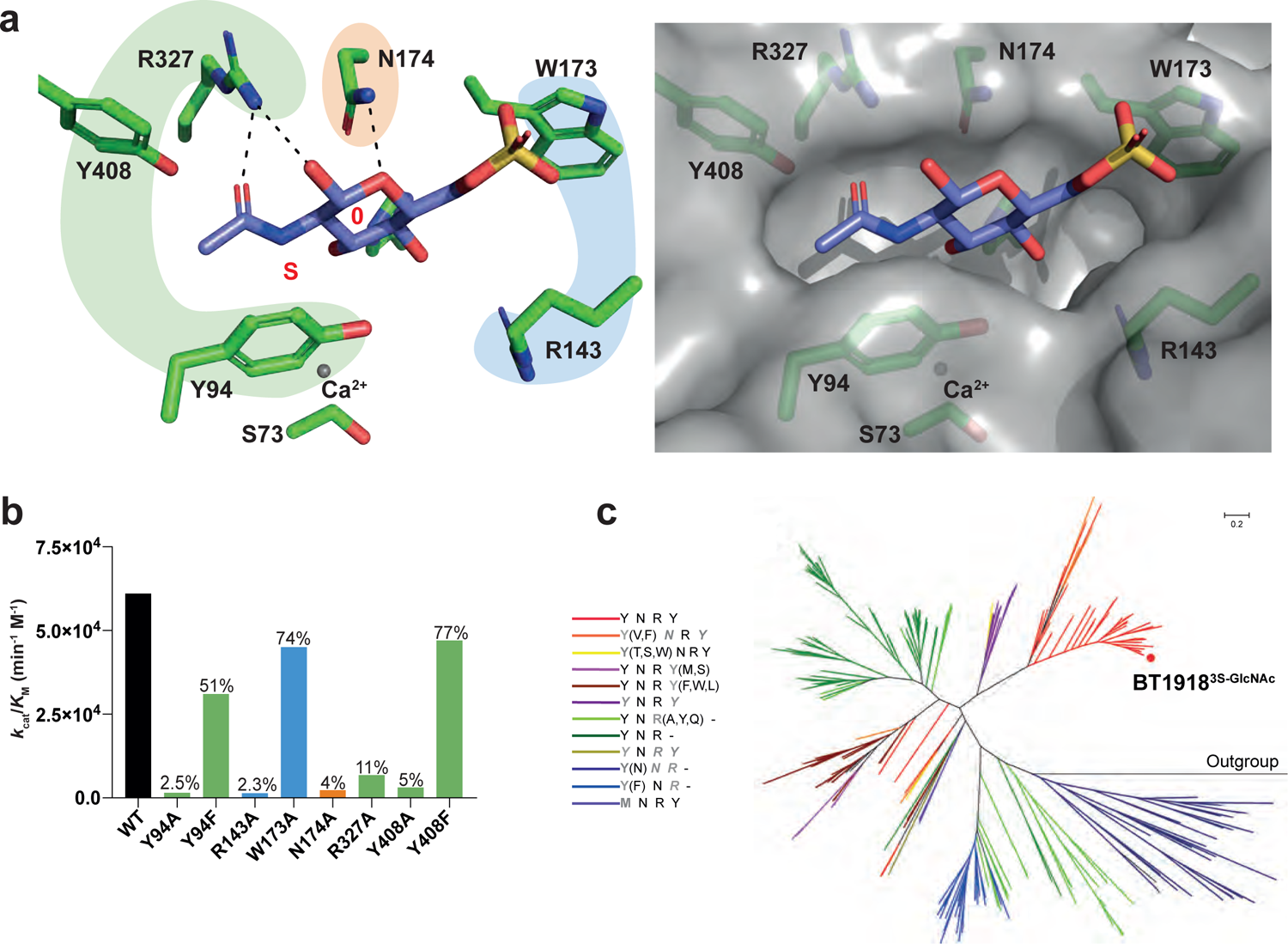
Structural details of the carbohydrate binding region of BT19183S-GlcNAc a, Stick representation of the structural data showing the carbohydrate binding interactions of the 0 subsite of BT1918^3S-GlcNAc^ (left panel) and surface representation of the structural data for BT1918^3S-GlcNAc^ showing the 0 subsite pocket (right panel). Highlighted in green are the residues interacting with the N-acetyl, in blue are the sulfate flanking residues, and orange indicates sugar ring only interactions. **b,** Catalytic effects of alanine scanning on BT1918^3S-GlcNAc^. Percentages above the bar indicate relative activity to wildtype (WT). **c,** Radial version of the phylogenetic tree of representative sulfatases from subfamily S1_46. For clarity all labels and sequence accession codes have been omitted. The annotations next to the colour code concern the presence or absence of conservation of the residues crucial in substrate recognition by BT1918^3S-GlcNAc^ (acc-code Q8A6G6) and in the following order: Y94, N174, R327 and Y408. The residues are coloured as following: black means an equivalent residue is present; a grey and bold letter at any position means that the corresponding residue is replaced by that amino acid; a grey, bold and italic letter at any position means that the equivalent position is replaced by any type of amino acid; a bold grey letter followed by one-letter codes in parentheses indicates that the equivalent position can be substituted by any of those amino acids; the dash at the Y408-equivalent position indicates that no equivalent amino acid can be deduced from the multiple alignment. Branches having the same colour have the corresponding pattern in common. The red filled circle designates the sequence of the S1_46 sulfatase from B. thetaiotaomicron (See Figure S8 for full tree).

### Aromatic stacking is essential for S1_16 4S-Gal/GalNAc sulfatases of the HGM

BT3057^4S-Gal/GalNAc^ and BT3796^4S-Gal/GalNAc^ are both 4S-D-galactose/*N*-acetylgalactosamine (4S-Gal/GalNAc) sulfatases, displaying a mix of monomeric and dimeric species, likely a consequence of heterologous expression, (**Figure S3 and Figure S4a,b,c**), and appear to bind Gal/GalNAc in similar manners (**Figure 4a,b**). In the 0 subsite of both sulfatases a critical aromatic residue, W109, stacks against the α face of the sugar ring whilst *O*3 is coordinated via Nε2 of H423, in BT3057^4S-Gal/GalNAc^, and the indole nitrogen of W431, in BT3796^4S-Gal/GalNAc^, (**Figure 4a, b**). In both enzymes mutation of W109 to Ala caused loss of any quantifiable activity but some trace activity was observed qualitatively, after extended incubations (**Table S1** and **Figure S4d,e**). The mutation of the residue coordinating *O*3 causes ∼10 and ∼50 fold reduction in activity for BT3057^4S-Gal/GalNAc^ and BT3796^4S-Gal/GalNAc^, respectively (**Table S1** and **Figure S4d,e**). In BT3057^4S-Gal/GalNAc^ an additional interaction is made via Nδ1 of H182 to *O*6 and mutation to Ala causes ∼20 fold reduction in activity (**Figure 4a** and **Table S1**). In BT3796^4S-Gal/GalNAc^ a glycine resides in place of H182 but a Trp residue, W332, lies opposite the position (**Figure 4b**). It is possible that both H182 (BT3057^4S-Gal/GalNAc^) and W332 (BT3796^4S-Gal/GalNAc^) residues could also contribute to the binding of a +1 sugar, through a stacking interaction, should such a substrate encounter the enzymes (**Figure 4a, b**). Interestingly, despite its critical nature, W109 is only observed in 37 % of analysed S1_16 sequences. However, this residue is found in 84 % of the sequences from organisms residing in the human gut, but drops to 17 % for sequences from a marine or aquatic environments where Val is most frequently observed (**Figure 4c and Table S3**). The aromatics Phe and Tyr, replace W109 at a frequency of ∼8 % and ∼6 %, respectively, and could theoretically perform the same stacking role as W109 (**Figure 4c and Table S3**). Bioinformatic analysis shows that 72 % of the total S1_16 sequences were from a marine or aquatic environment with terrestrial and human sources making up most of the remainder in roughly equal proportion (**Table S3**). These data suggest that only the subset of S1_16 sulfatases possessing an equivalent residue to W109 will be active as 4S-Gal/GalNAc sulfatases and that this activity is mostly restricted to human gut bacteria; S1_16 sulfatases from marine environments likely target a different, but potentially somewhat analogous, sulfated linkage.

**Figure 4.**
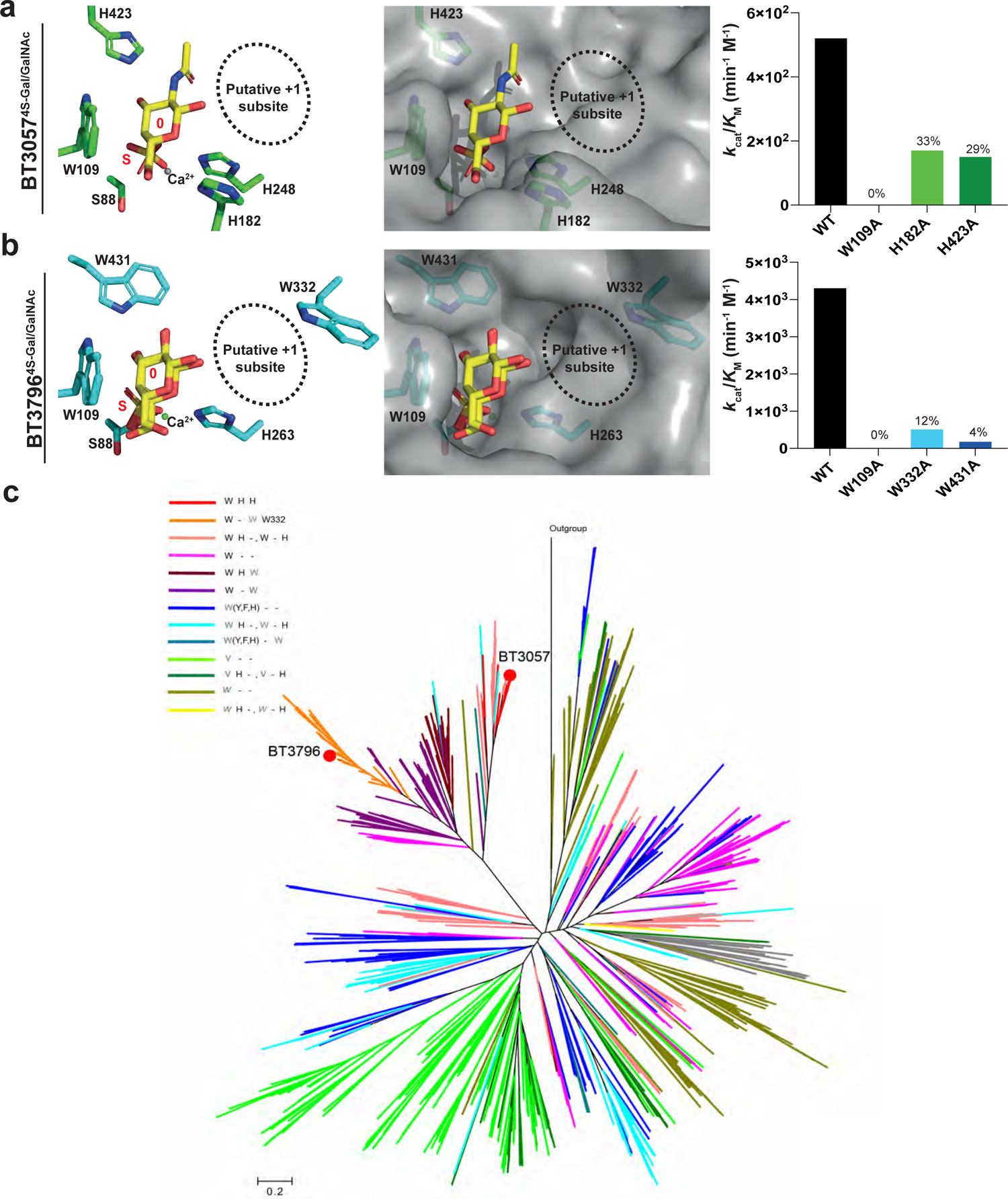
Structural details of the carbohydrate binding region of BT30574S Gal/GalNAc and BT37964S Gal/ GalNAc. Stick (left) and surface (middle) representation showing the carbohydrate binding interactions of the 0 subsite of S1_16 sulfatases: **a,** BT3057^4S-Gal/GalNAc^ and **b,** BT3796^4S-Gal/GalNAc^. The respective right panel show the effects of alanine scanning on these sulfatase activities with the percentages above the bar indicating the relative activity to wildtype (WT). **c,** Radial phylogenetic tree of representative sulfatases from subfamily S1_16. For clarity all labels and sequence accession codes have been omitted. The annotations next to the colour code concern the presence or absence of conservation of the critical residues in substrate recognition by BT3057^4S-Gal/GalNAc^ (acc-code Q8A397) in the order: W109, H182 and H423. Sequences coded by orange branches contain an additional W332 present in BT3796^4S-Gal/GalNAc^ (acc-code Q8A171) but absent in other sequences. For simplification the residue numbers have been omitted, except for W332. The residues are coloured as following: black means an equivalent amino acid is present; a grey and bold letter at any position means that the corresponding residue is replaced by that amino acid; a grey and italic letter at any position means that the equivalent position is replaced by any type of amino acid; a bold grey letter followed by one-letter codes in parentheses indicates that the equivalent position can be substituted by any of those amino acids; the dash at the H-equivalent position indicates that no equivalent amino acid can be deduced from the multiple alignment. When two patterns are indicated separated by a comma (i.e. W - H, W H -) both have been attributed the same colour code. Branches having the same colour have the corresponding pattern in common. Red filled circles designate sequences of S1_16 sulfatases from B. thetaiotaomicron (See Figure S9 for full tree).

### Defining the conserved and variable features driving 6S-Gal/GalNAc recognition

The 4 S1_15 enzymes characterised here are all monomeric (**Figure S5a**) and exclusively target 6S-D-galacto configured substrates^10^ through a conserved mechanism that was observed previously for BT3333^6S-GalNAc^, a sulfatase that specifically cleaves 6S-GalNAc^9^. These conserved recognition features are D176, R177 and H227 in BT4631^6S-Gal/GalNAc^, D170, R171, and H220 in BT1624^6S-Gal/GalNAc^, and D162, R163 and H212 in BT3109^6S-Gal^ (**Figure 5a, b**). In all three enzymes the His residue co-ordinates with *O*3 via Nε2, whilst Asp and Arg hydrogen bond to *O*4 through Oδ2 and Nη1, respectively (**Figure 5a**). The His residue resides in a small motif (HDQSIV) present in 56 % of all sequences but, within the motif, the His exists at a 99 % frequency. The Asp and Arg are found within the conserved motif FIMAATGDRVP but Asp and Arg are only found at frequencies of 74 % and 56 %, respectively (**Extended data 1** and **Table S4**). Overall this means the ‘recognition triad’ is conserved in 56 % (523) of all sequences analysed, suggesting that at least half of the members of this subfamily target galacto configured carbohydrates. Analysis of the environment from which the S1_15 sequences were derived reveal that ∼65 % of all S1_15 sequences are from a marine environment and that nearly 70 % of these lack the galactose recognition triad (**Table S4**). By contrast, 83 %, 80% and 95 % of sequences from human, animal, and terrestrial sources, respectively, contain the galactose recognition triad (**Table S4**). This suggests that S1_15 subfamily members target at least two types of sulfated substrates; galacto configured monosaccharides and an unknown sulfate conjugate which is enriched in the marine environment. Alanine scanning was carried out on the 0 subsite of BT1624^6S-Gal/GalNAc^ to further inform its carbohydrate binding interactions but no kinetic data could be obtained. Qualitative analysis by TLC and HPAEC however, revealed that the mutants were active but that against 6S-Gal mutation of any residues in the ‘recognition triad’ resulted in either complete loss, or trace levels of activity, whilst against 6S-GalNAc only the D170A mutant resulted in complete loss of activity (**Figure 5c and S6a**). DSF analysis indicates that all proteins were folded but were less stable, with exception of R171A which showed an increase in stability (**Figure S6b**). As expected, the mutants bound Gal more weakly than the wildtype BT1624^6S-Gal/GalNAc^ (**Figure S6c**).

**Figure 5.**
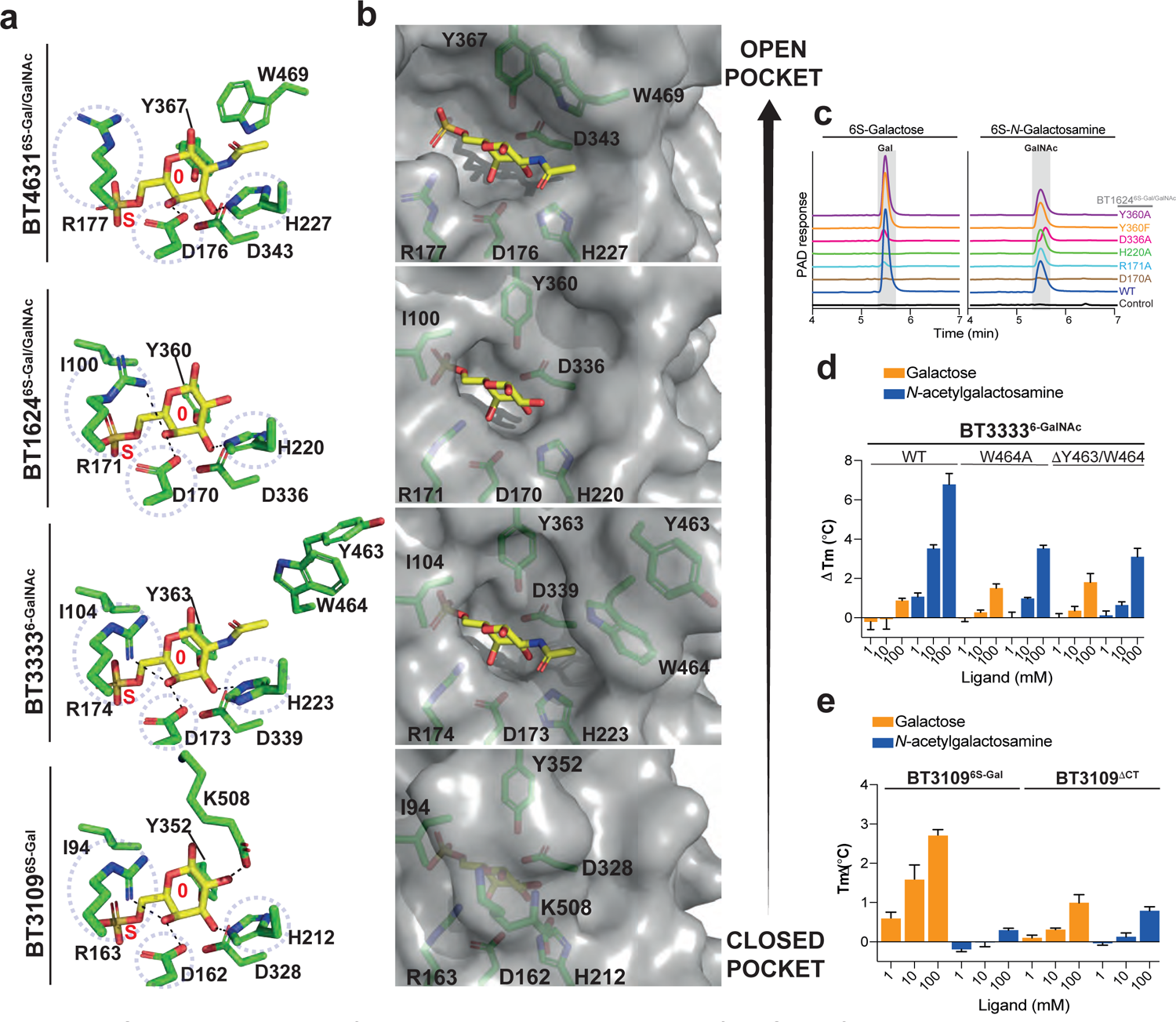
Structural variations of the carbohydrate binding region of the S1_15 family a, Stick and **b,** Surface representation showing the carbohydrate binding interactions of the 0 subsite of BT4631^6S-Gal/GalNAc^, BT1624^6S-Gal/GalNAc^, BT3333^6S-GalNAc^, and BT3109^6S-Gal^ (from top to bottom). **c,** High pressure anion _exchange_ chromatography (HPAEC) of wildtype BT1624^6S-Gal/GalNAc^ wildtype (WT) and its mutants. The produced product is highlighted by a grey box. HPAEC reactions utilised 6 mM substrate and 5 M enzyme, with 3 mM HEPES, 45 mM NaCl, and 5 mM CaCl2 over a 48 h period at 37°C. **d,** DSF analysis of the effect of mutating the GalNAc specificity features of BT3333^6S-GalNAc^. **e,** DSF analysis of the effect of mutating the C-terminal extension (VEEEPLK) which drives specificity towards Gal in BT3109^6S-Gal^. 100 mM BTP pH 7.0 with 150 mM NaCl was used in all DSF experiments. Error bars represent s.e.m..

A key difference between the structures of the 4 S1_15 enzymes is in the openness of their respective subsites (**Figure 5b**). BT4631^6S-Gal/GalNAc^ displays a more open S subsite with the C6 sulfate solvent exposed and the 0 subsite may allow the accommodation of additional groups. It is likely that true substrate targeted by this sulfatase remains to be discovered. In the other 3 structures, an Ile is found over the S subsite and the sulfate group is buried in a deep pocket, whilst further differences in the residues interacting with C2 substituents drive variation in substrate preference. BT1624^6S-Gal/GalNAc^ forms few or no interactions with the *N*-acetyl group of GalNAc or the *O*2 of Gal and has a more open pocket than BT3333^6S-GalNAc^ (**Figure 5a, b**). Indeed, BT1624^6S-Gal/GalNAc^ displays equal affinity, and activity, on both 6S-Gal and 6S-GalNAc^10^ (**Table S5**). By contrast, in BT3333^6S-GalNAc^ W464 is positioned to interact with the *N*-acetyl group of GalNAc through a stacking interaction with Y463 (**Figure 5a**). This leads to the enzyme having both a greater affinity and activity on 6S-GalNAc than BT1624^6S-Gal/GalNAc^, and no detectable activity on 6S-Gal (**Table S5**)^10^. Mutation of W464 to Ala, or deletion of Y463/W464, significantly reduces the preference of BT3333^6S-GalNAc^ for GalNAc, whilst slightly increasing its interaction with Gal (**Figure 5d**). It is interesting to note that W464 is also present in BT4631^6S-Gal/GalNAc^ (W469) but the Y463 is replaced by T464, a less bulky amino acid (**Figure 5a**). In the absence of the positioning aromatic, W469 flips down and is unable to interact with the *N*-acetyl of the 0 subsite sugar. Additional bioinformatic analysis of S1_15 enzymes located within *Bacteroides* PULs targeting chondroitin sulfate (CS), showed a retention of W464. However, Y463 is only partially retained, being mainly replaced by the aromatic residues Phe and His, which are also capable of stacking against W464 thus, preserving its functionality to interact with the *N*-acetyl group of GalNAc (**Extended data 2**). Together these data suggest that the sulfatases located within CS PULs have evolved to specifically target 6S-GalNAc. In comparison to the other 3 S1_15 enzymes, BT3109^6S-Gal^ forms an additional interaction with the O2 of Gal via the carboxy terminus of K508 (**Figure 5a, b**). This interaction appears to exclude the *N*-acetyl group from binding explaining the strong preference for 6S-Gal over 6S-GalNAc substrates, as indicated by activity and affinity studies (**Figure 5e** and **Table S5**). To further confirm this result, we generated a mutant (BT3109^ΔCT^) where we removed the carboxy terminus region (VEEEPLK) of BT3109^6S-Gal^. The BT3109^ΔCT^ mutant did not show a preference for Gal and bound both Gal and GalNAc with a similar affinity (**Figure 5e**). This indicates that the terminal region of BT3109^6S-Gal^ tailors substrate specificity towards Gal. Interestingly, the VEEEPLK sequence is only found in 50 other sequences analysed and is found almost exclusively in S1_15 sulfatases from Bacteroidetes inhabiting marine environments (**Table S6**).

### Comparison of the S1_11 sulfatases BT3177^6S-GlcNAc^ and BT4656^6S-GlcNAc/GlcNS^

BT3177^6S-GlcNAc^ and BT4656^6S-GlcNAc/GlcNS^ are both monomeric (**Figure S5b**) and utilise D361/D385 and R363/R387 to coordinate the *O*4 through Oδ2, and Nη1, respectively, and H445/H471 to coordinate *O*3 via Nε2 (**Figure 6**). The key role of these residues in substrate recognition was confirmed by alanine scanning of the 0 subsite of BT3177^6S-GlcNAc^ (**Table S5**). The D361A mutation causes complete loss of activity whilst R363A and H445A cause ∼15 and ∼1500-fold loss in catalytic activity, respectively (**Table S5**). The Asp, Arg, and His are found within conserved motifs and are present in 91 % of all 955 representative sequences analysed, with Asp and Arg even higher at 98 and 99% (**Extended data 3** and **Table S7**). This suggests that the S1_11 subfamily may exclusively target 6S-D-gluco configured carbohydrates.

**Figure 6.**
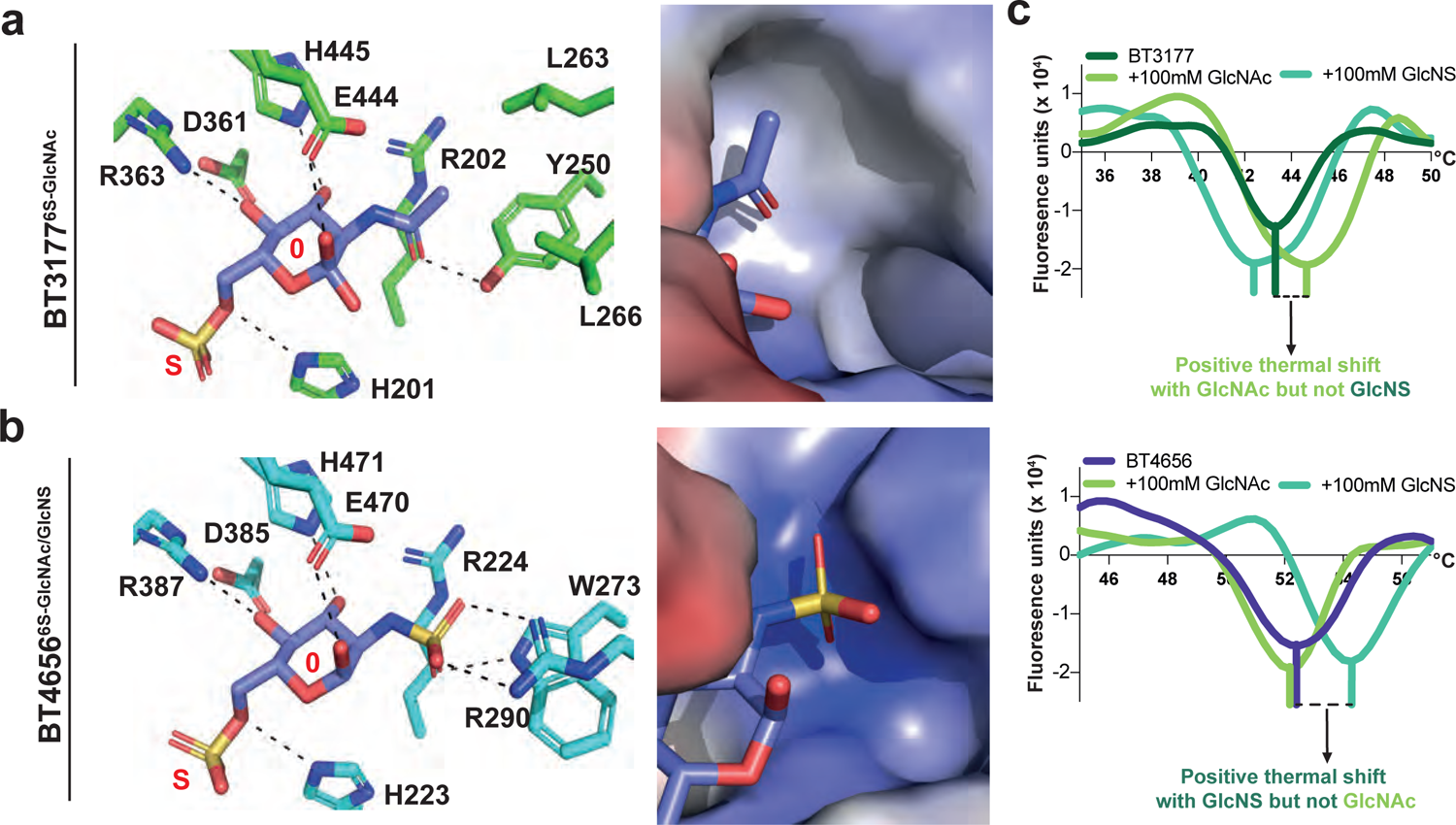
Structural variations of the carbohydrate binding region of the S1_11 family a, Carbohydrate interactions of BT3177^6S-GlcNAc^. **b,** Carbohydrate interactions of BT4656^6S-GlcNAc/GlcNS^. In both panels the left is a stick representation of the carbohydrate interactions at S0, middle is a surface representation showing the local surface charge. **c,** DSF data showing the second derivative of the thermal melts with protein alone, 100 mM GlcNAc, and 100 mM GlcNS against 5 **u**M BT3177^6S-GlcNAc^ (top), and 5 μM of _BT4656_6S-GlcNAc/GlcNS _(bottom)._

BT3177^6S-GlcNAc^ and BT4656^6S-GlcNAc/GlcNS^ have no discernible differences in the residues recognising GlcNAc/GlcNS unless focus is directed toward the amino acids coordinating the *N*-acetyl group and the *N*-sulfate groups, respectively. This region shows high amino acid diversity and is absent in around half of the S1_11 sequences analysed (**Extended data 3** and **Table S7**). BT3177^6S-GlcNAc^ hydrogen bonds to the carbonyl of the *N*-acetyl group through the phenol OH of Y250, and the overall environment has a hydrophobic character having L263 and L266 sitting above Y250 (**Figure 6a**). Mutation of Y250 to Ala caused a ∼10-fold loss in activity whilst mutation to Phe had no effect demonstrating that only the hydrophobic character of the aromatic phenyl ring is important (**Table S5**). By contrast, BT4656^6S-GlcNAc/GlcNS^ has W273 and R290 situated in this region, whilst L263 is absent leaving a more open landscape (**Figure 6b**). R290 sits above W273 with the two residues interacting through cation-π interactions, positioning R290 to form a bidentate ionic interaction with the *N*-sulfate group, whilst the Nε1 of the Trp indole ring forms a hydrogen bond with the third oxygen of the sulfate. This gives this area of the protein a far more basic charge as compared to BT3177^6S-GlcNAc^ (**Figure 6**). Despite the high variability of this region (**Extended data 3** and **Table S7**), homologues of BT4656^6S-GlcNAc/GlcNS^, harbouring W273/R290, are highly conserved in PULs targeting the glycans Hep/HS where 6S-GlcNS is enriched compared to other sulfated substrates (**Extended data 4**). Although, BT4656^6S-GlcNAc/GlcNS^ is active on 6S-GlcNAc, also found in Hep/HS, the presence of R290/W273 may provide enhanced activity towards 2*N*-sulfated 6S-GlcNAc compared to BT3177^6S-GlcNAc^. Indeed, although the *k*_cat_/*K*_M_ is similar for both substrates (**Table S5**), DSF analysis shows that BT4656^6S-GlcNAc/GlcNS^ has a greater affinity for GlcNS than GlcNAc, whilst the reciprocal is true for BT3177^6S-GlcNAc^ (**Figure 6b**). This result is consistent with BT3177^6S-GlcNAc^ being encoded in a PUL associated with mucin degradation^10^ and therefore more tailored towards 6S-GlcNAc, a sulfated linkage common in mucin *O*-glycans.

Analysis of the environment from which S1_11 sequences were isolated shows that around 50 % come from a marine origin whilst, S1_11 has a 2-3 fold enrichment in sequences (∼27% total) from a terrestrial/soil based environment when compared to the S1_15, S1_16, and S1_46 subfamilies (**Table S7**). Interestingly, despite 6S-GlcNAc being a common component of host glycans this subfamily contains relatively few sequences (15 %) from the host environment. Conversely, 6S-gluco configured substrates are not well associated with marine environments. This observation suggests that marine S1_11 sulfatases utilise the invariant recognition triad to target a different substrate to the S1_11 sulfatase characterised here or to target an unknown substrate rich in 6S-gluco configured sugars.

## Discussion

CAZymes are exquisitely specific enzymes, often distinguishing between single epimeric features to drive their specificity. S1 carbohydrate sulfatases are no exception to this. S1_11 and S1_15 subfamilies utilise an identical ‘recognition triad’ but specificity for GlcNAc and GalNAc (epimeric at *O*4) is achieved by the residues coming from the C-terminus in S1_11 and the N-terminus in S1_15, meaning no interactions are spatially conserved and thus recognition is specific (see supplemental discussion, **Figure S7**, and **Table S8** for further comparison). The finer details of recognition, however, are ‘hidden’ in the non-conserved, and highly variable, area of the proteins. In these regions BT3177^6S-GlcNAc^ and BT3333^6S-GalNAc^ have both evolved aromatic residues capable of interacting with the *N*-acetyl group, of their respective substrates. BT3177^6S-GlcNAc^ resides in a PUL targeting mucin *O*-glycans where 6S-GlcNAc is a common component, whilst BT3333^6S-GalNAc^ resides in a PUL targeting CS where 6S-GalNAc can comprise up to 50 % of the polymer (**Figure 1**). By contrast BT1624^6S-Gal/GalNAc^, also in a PUL targeting mucin *O*-glycans, shows no tailored adaptation in this region, and as such has similar activity on both 6S-Gal and 6S-GalNAc. Finally, BT4656^6S-GlcNAc/GlcNS^, which resides in a PUL targeting the GAGs Hep and HS, has a positively charged nature in the equivalent area. This provides a stronger interaction with the doubly sulfated 6S-GlcNS, a component found exclusively in Hep and HS. Interestingly, GAGs have been shown to be high priority substrates for several *Bacteroides* species^17^ and the enhanced activities of sulfatases targeting host glycans may bestow a critical advantage on these substrates within the competitive gut environment.

BT3109^6S-Gal^ shows an unusual C-terminal feature where the C-terminal carboxyl group caps the active site excluding the ability to recognize GalNAc. BT3109^6S-Gal^ resides in a PUL of unknown function but also contains a GH2 and GH43_31, both of which can target Gal in the pyranose and furanose form, respectively. Therefore, combined with the knowledge that BT3109^6S-Gal^ orthologues are most commonly found in marine environments, we suggest that this PUL may target a marine polysaccharide containing 6S-Gal, an example of which being λ-carrageenan or potentially a more enigmatic marine glycan not yet identified^25^. Indeed, the ability of the human gut microbiota to metabolise marine glycans was recently revealed to be more extensive than previously thought^26^ suggesting these glycans may make a significant contribution to the colonic ecosystem.

The S1_46 subfamily sequences from both Firmicutes and Bacteroidetes inhabiting the human gastrointestinal tract require the *N*-acetyl group for activity. All the key residues are invariant with the exception of Y408 that is preserved in Bacteroidetes but is mostly absent in gut Firmicutes. Intriguingly, these recognition features were also found in several pathogens suggesting that S1_46 sulfatases could be important enzymes for accessing host glycans in these species. For S1_46 sequences from the marine environment significant variability is seen in the key residues recognising the *N*-acetyl group suggesting this not a feature encountered in their substrate. The sequences from the gastrointestinal tract most closely related to those from the marine environment also display similar mutations thus, despite their differing environments these sequences may target a similar, as yet unidentified, substrate to their marine counterparts. This observation further strengthens support for recent data highlighting marine glycan metabolism within the HGM.

The enrichment of sequences from a terrestrial/soil based environment in S1_11 is intriguing due to the presence of 6S-GlcNAc in nodulation (NOD) factors; bacterially produced lipooligosaccharides essential for bacteria to invade plant roots and establish nitrogen fixing nodules^27^. Indeed, several bacterial S1_11 sequences were isolated from rhizosphere communities and root nodules suggesting that S1_11 sulfatases could be involved in NOD factor metabolism. Interestingly, S1_11 sequences within the fungal phylum Ascomyota are well represented in plant pathogens. This alludes to a potential dichotomy where plant associated bacterial species may utilise S1_11 sulfatases to de-sulfate NOD factors in a positive manner, whilst S1_11 sequences in fungal Ascoymota may act as virulence factors, potentially by hijacking the NOD factor system. It should be noted however that the S1_11 sulfatases here are exo-acting, from the non-reducing end, and that NOD factors are sulfated at the reducing end of their chito-oligosaccharide component. This does not readily lend itself to a mechanism where S1_11 sulfatases act directly on intact NOD factors but these sulfatases may instead be involved in their downstream catabolism.

## Conclusion

S1 carbohydrate sulfatases are exquisitely tailored enzymes deriving their binding energy and specificity from the nature of the glycan to which the target sulfate is appended. This makes these enzymes excellent targets for small molecule intervention to modify their function, develop tools to probe sulfated glycan metabolism, and disease intervention where sulfated glycan metabolism is perturbed.

## Materials and methods

### Recombinant Protein Production

Genes were amplified by PCR using the appropriate primers and the amplified DNA cloned into pET28b using NheI/XhoI restriction sites generating constructs with N-terminal His_6_ tags (Supplementary Table 11). Recombinant genes were expressed in *Escherichia coli* strains BL21 (DE3) or TUNER (Novagen), containing the appropriate recombinant plasmid, and cultured to mid-exponential phase in LB supplemented with 50 μg/mL kanamycin at 37 °C and 180 rpm. Cells were then cooled to 16 °C, and recombinant gene expression was induced by the addition of 0.1 mM isopropyl β-D-1-thiogalactopyranoside; cells were cultured for another 16 h at 16 °C and 180 rpm. The cells were then centrifuged at 5,000 × g and resuspended in 20 mM Hepes, pH 7.4, with 500 mM NaCl before being sonicated on ice. Recombinant protein was then purified by immobilized metal ion affinity chromatography using a cobalt-based matrix (Talon, Clontech) and eluted with 100 mM imidazole. For the proteins selected for structural studies, another step of size exclusion chromatography was performed using a Superdex 16/60 S75 or S200 column (GE Healthcare), with 10 mM HEPES, pH 7.5, and 150 mM NaCl as the eluent, and they were judged to be ≥95% pure by SDS-PAGE. Protein concentrations were determined by measuring absorbance at 280 nm using the molar extinction coefficient calculated by ProtParam on the ExPasy server (web.expasy.org/protparam/).

### Site-Directed Mutagenesis

Site-directed mutagenesis was conducted using the PCR-based QuikChange kit (Stratagene) according to the manufacturer’s instructions using the appropriate plasmid as the template and appropriate primer pairs (**Table S9**).

### Microfuidics de-sulfation assays

Sulfated carbohydrates were labelled at their reducing end with BODIPY which has a maximal emission absorbance of ∼503nm, which can be detected by the EZ Reader via LED-induced fluorescence. Non-radioactive microfluidic mobility shift carbohydrate sulfation assays were optimised in solution with a 12-sipper chip coated with CR8 reagent and a PerkinElmer EZ Reader II system using EDTA-based separation buffer and real-time kinetic evaluation of substrate de-sulfation. Pressure and voltage settings were adjusted manually (1.8 psi, upstream voltage: 2250 V, downstream voltage: 500 V) to afford optimal separation of the sulfated and unsulfated product with a sample (sip) time of 0.2 s, and total assay times appropriate for the experiment. Individual de-sulfation assays were carried out at 28°C and assembled in a 384-well plate in a volume of 80 μl in the presence of substrate concentrations between 0.5 and 20 μM with 100 mM Bis-Tris-Propane or Tris, depending on the pH required, 150 mM NaCl, 0.02% (v/v) Brij-35 and 5 mM CaCl_2_. The degree of de-sulfation was directly calculated using the EZ Reader software by measuring the sulfated carbohydrate: unsulfated carbohydrate ratio at each time-point. The activity of sulfatase enzymes was quantified in ‘kinetic mode’ by monitoring the amount of unsulfated glycan generated over the assay time, relative to control assay with no enzyme; with sulfate loss limited to ∼20% to prevent of substrate and to ensure assay linearity. *k*_cat_/*K*_M_ values, using the equation V_0_=(V_max_/*K*_M_)[S], were determined by linear regression analysis with GraphPad Prism software. Substrate concentrations were halved and doubled to check linearity of the rates ensuring substrate concentrations were significantly <*K*_M_.

### HPAEC and TLC sulfatase enzymatic assays

For reactions analysed by thin layer chromatography (TLC) 2 μL of each sample was spotted onto silica plates and resolved in butanol:acetic acid:water (2:1:1) running buffer. The TLC plates were dried, and the sugars were visualized using diphenylamine stain (1 ml of 37.5% HCl, 2 ml of aniline, 10 ml of 85% H3PO3, 100 ml of ethyl acetate and 2 g diphenylamine) and heated at 450°C for 2-5 min with a heat gun. Where possible, the enzymatic activity was confirmed by high-performance anionic exchange chromatography (HPAEC) with pulsed amperometric detection using standard methodology. The sugars (reaction products) were bound to a Dionex CarboPac PA200 column and eluted with an isocratic flow of 80 mM NaOH for 15 min, the column was then cleaned with 500 mM NaOH for 10 min before being ran back into 80 mM NaOH at a flow rate of 0.25 ml min^-1^ before injection of the next sample. The reaction products were identified using the appropriated standards.

### Differential scanning fluorimetry

Thermal shift/stability assays (TSAs) were performed using a StepOnePlus Real-Time PCR machine (LifeTechnologies) and SYPRO-Orange dye (emission maximum 570 nm, Invitrogen) as previously described^28^ with thermal ramping between 20 and 95°C in 0.3°C step intervals per data point to induce denaturation in the presence or absence of various carbohydrates as appropriate to the sulfatase being analysed. The melting temperature (Tm) corresponding to the midpoint for the protein unfolding transition was calculated by fitting the sigmoidal melt curve to the Boltzmann equation using GraphPad Prism, with R^2^ values of >0.99. Data points after the fluorescence intensity maximum were excluded from the fitting. Changes in the unfolding transition temperature compared with the control curve (ΔT_m_) were calculated for each ligand. A positive ΔT_m_ value indicates that the ligand stabilises the protein from thermal denaturation, and confirms binding to the protein. All TSA experiments were conducted using a final protein concentration of 5μM in 100 mM Bis-Tris-Propane (BTP), pH 7.0, and 150 mM NaCl supplemented with the appropriate ligand. Three independent assays were performed for each protein and protein ligand combination (**Table S10 and 11)**.

### Glycan labelling

Sulfated saccharide samples were labelled according to a modification of the method by Das *et al*., reporting the formation of N-glycosyl amines for 4,6-O-benzilidene protected D-gluopyranose monosaccharides with aromatic amines^29^. Briefly, the lyophilised sugar (1 mg) was dissolved in anhydrous methanol (0.50 mL, Sigma-Aldrich) in a 1.5 mL screw-top PTFE microcentrifuge tube and BODIPY-FL hydrazide (4,4-difluoro-5,7-dimethyl-4-bora-3a,4a-diaza-*s*-indacene-3-propionic acid, hydrazide, 0.1 mg, ThermoFischer, λ_ex./em._ 493/503, ε 80,000 M^-1^cm^-1^) was added and the mixture vortexed (1 min), then reacted (65 °C, 24 h) in darkness. The products were then cooled and a portion purified by TLC on silica coated aluminium plates (silica gel 60, Sigma-Aldrich, Millipore) developed with methanol or 1:1 v/v ethyl acetate/methanol. The unreacted BODIPY-FL label (orange on the TLC plate) was identified by reference to a lane containing the starting material (BODIPY-FL hydrazide), allowing differentiation from the putative labelled product (also orange). This latter band (which can run ahead or behind the label depending on the sugar; e.g. labelled GlcNAc 4S and 6S both run with R_f_ 0.84 compared to label, R_f_ 0.70; others, such as labelled GalNAc4S or 6S require, 1:1 v/v ethyl acetate/methanol and the product runs behind the label on TLC) was scraped from the plates and extracted in fresh methanol (2 x 0.5 mL), spun (benchtop centrifuge, 3 minutes), the supernatant recovered and dried (rotary evaporator) to afford the fluorescent, coloured product (bright green in aqueous solution), which was then employed in subsequent experiments.

### Isothermal Calorimetry (ITC)

The affinity of BT3333^6S-GalNAc^ and BT4656^6S-GlcNAc/GlcNS^ against 6S-GalNAc and 6S-GlcNAc, respectively, was quantified by ITC using a Microcal ITC^200^ calorimeter. The protein samples (70 μM for BT3333^6S-GalNAc^ and 60 μM for BT4656^6S-GlcNAc/GlcNS^), stirred at 400 rpm in a 0.2-mL reaction cell, was injected with 18 2 μL aliquots of ligand, preceded by 1 injection of 0.2 μL with a delay of 180 seconds between injections (0.8 mM 6S-GalNAc was used for BT3333^6S-GalNAc^ and 0.4 mM 6S-GlcNAc for BT4656^6S-GlcNAc/GlcNS^). Titrations were carried out in 50 mM Tris-HCl buffer, pH 8.0, at 25 °C. Integrated binding heats minus dilution heat controls were fit to a single set of sites binding model to derive *K*_A_, ΔH, and n (number of binding sites on each molecule of protein) using Microcal Origin v7.0.

### Static light scattering determination of molecular weight

Molecular weights were determined using an Agilent Multi-Detector System calibrated with bovine serum albumin. Proteins were separated by size exclusion chromatography using an Agilent BioSEC Advance 300 Å, 4.6 x 300 mm or GE Superdex 200 10 300 columns equilibrated with 20 mM tris(hydroxymethyl)aminomethane-HCl pH 7.4, 150 mM NaCl buffer. Light scattering data was collected at 90° and refractive index used to calculate absolute molecular weight.

### Crystallisation of carbohydrate sulfatases

After purification, all proteins were concentrated in centrifugal concentrator with a molecular weight cutoff of 30 KDa in the size exclusion chromatography buffer. Sparse matrix screens were set up in 96-well sitting drop SPT Labtech plates plates (400-nL drops). Initial hits crystals for all proteins were obtained between 20 and 35 mg/mL supplemented with between 10 and 30 mM ligand unless otherwise stated. For all sulfatase the wildtype *B. theta* variants were used, having a Ser at the catalytic formylglycine position. BT1624^6S-Gal/GalNAc^ with 6S-GalNAc crystallised in 20 % Polyethyleneglycol (PEG) 6000, 0.2 M ammonium chloride and 0.1 M sodium acetate pH 5.5. BT1918^3S-GlcNAc^ with 6S-GlcNAc crystallised in 45 % Methylpentanediol (MPD) 0.2 M CaCl_2_ and 0.1 M Bis-Tris pH 5.5. BT3057^4S-Gal/GalNAc^ with 4S-GalNAc crystallised in 20 % PEG 3350 and 0.2 M sodium nitrate. BT3109^6S-Gal^ with 6S-Gal crystallised in 30 % PEG 4000, 0.2 M ammonium acetate and sodium citrate pH 5.6. BT3177^6S-GlcNAc^ with 6S-GlcNAc crystallised in 50 % precipitant mix 1 (40% v/v PEG 500 MME; 20 % w/v PEG 20000), 0.1 M carboxcylic acids (0.2 M Sodium formate; 0.2 M Ammonium acetate; 0.2 M Sodium citrate tribasic dihydrate; 0.2 M Sodium potassium tartrate tetrahydrate; 0.2 M Sodium oxamate) and 0.1 M buffer system 3 pH 8.5 (Tris (base); BICINE). BT3796^4S-Gal/GalNAc^ with 4S-GalNAc crystallised in 20 % PEG 6000, 0.2 M magnesium chloride and 0.1 MES pH 6.0. BT4631^6S-Gal/GalNAc^ was crystallised at 80 mg/ml with 10 mM 6S-Gal in 20 % PEG 10000 with 0.1 M Bicine pH 8.5. All crystals were cryo-cooled with the addition of the ligand they were crystallised with. 20% PEG 400 was used as the cryoprotectant for BT1624^6S-Gal/GalNAc^, BT3057^4S-Gal/GalNAc^, and BT3109^6S-Gal^ and 100 % paratone-N oil for BT3796^4S-Gal/GalNAc^. PEG 200 was used as cryoprotectant for BT4631^6S-Gal/GalNAc^ crystals. No cryoprotectant was added to BT1918^3S-GlcNAc^ or BT3177^6S-GlcNAc^ crystals as the crystallisation condition afforded sufficient cryoprotection. Data were collected at Diamond Light Source (Oxford) on beamlines I03, I04, I04-1 and I24, and SOLEIL on the PROXIMA_1 beamline at 100 K. The data were integrated with XDS^30^, or Xia2 3di or 3dii, and scaled with Aimless^31, 32^. Five percent of observations were randomly selected for the R_free_ set. The phase problem was solved by molecular replacement using the automated molecular replacement server Balbes^33^ for all proteins except BT3109^6S-Gal^ and BT4631^6S-Gal/GalNAc^ which were solved using Phaser^34^ and BT1624^6S-Gal/GalNAc^ as the search model after preparation with sculptor. Models underwent recursive cycles of model building in Coot^35^ and refinement cycles in Refmac5^36^. Where necessary ligand restraint and coordinates were generated with Jligand^37^. The models were validated using Coot and MolProbity^38^. Structural Figures were made using Pymol (The PyMOL Molecular graphics system, Version 2.0 Schrodinger, LLC.) and all other programs used were from the CCP4 suite^39, 40^. The data processing and refinement statistics are reported in **Table S12 and S13**.

### Global phylogenetic trees of S1_11, S1_15, S1_16, S1_46 sequences

On the basis of the taxonomic diversity, to avoid identical sequences we selected a representative number of sequences within each subfamily: a) for S1_11, 955 sequences were selected among the 2177 sequences present in the subfamily S1_11 from the SulfAtlas database and 411 positions were used for phylogeny; b) for S1_15, 920 sequences were selected among the 1906 sequences present in SulfAtlas and 365 positions were used for phylogeny; c) for S1_16, 800 sequences were selected among the 1361 sequences present in Sulfatlas and 342 positions were used for phylogeny; and d) for S1_46, 349 out of the total 574 sequences present in Sulfatlas were selected, 401 positions were used for phylogeny. In each case, the sequences were aligned by MAFFT v.7^41^ using L-INS-i algorithm. The multiple sequence alignments were visualized by Jalview software v.11.0^42^, non-aligned regions were removed, and the above listed respective numbers of positions were used for the phylogeny. Phylogeny was made using RAxML v. 8.2.4^43^. The phylogenetic tree was build with the Maximum Likelihood method^44^ and the LG matrix as evolutive model^45^ using a discrete Gamma distribution to model evolutionary rate differences among sites (4 categories). The rate variation model allowed for some sites to be evolutionarily invariable. The reliability of the trees was tested by bootstrap analysis using 1000 resamplings of the dataset^46^. In all cases, fifteen S1_0 sequences from the sulfAtlas database were used as outgroup.

### Data availability statement

Source Data for all experiments, along with corresponding statistical test values, where appropriate, are provided within the paper and in Supplementary information. The crystal structure dataset generates have been deposit in the in the Protein Data Bank (PDB) under the following accession numbers: 7OZ8, 7OZ9, 7OZA, 7OZE, 7OZC, 7P26, and 7P24.

### Code availability statement

No new codes were developed or compiled in this study

## Supporting information

Supplemental discussion and figures

S1_46 sequences detailing protein accession number, species, phylum, and the environmental context of each sequence.

S1_16 sequences detailing protein accession number, species, phylum, and the environmental context of each sequence.

S1_15 sequences detailing protein accession number, species, phylum, and the environmental context of each sequence.

S1_11 sequences detailing protein accession number, species, phylum, and the environmental context of each sequence.

## Acknowledgements

This project has received funding from the European Union’s Horizon 2020 research and innovation programme under the Marie Skłodowska-Curie grant agreement N° 748336, the European Research Council ERC (694181), The Knut and Alice Wallenberg Foundation (2017.0028), Swedish Research Council (2017-00958), Wilhelm och Martina Lundgrens Vetenskapsfond (2020.3597, awarded to ASL) and the Academy of Medical Sciences/Wellcome Trust through the Springboard Grant SBF005\1065 163470 awarded to AC. The authors acknowledge access to the SOLEIL and Diamond Light sources via both the University of Liverpool and Newcastle University BAGs (proposals mx21970 and mx18598, respectively). We thank the staff of DIAMOND, SOLEIL, and members of the Liverpool’s Molecular biophysics group for assistance with data collection. We are also grateful for Dr. Erwan Corre’s help regarding bioinformatics analyses (ABIMS platform, Station Biologique de Roscoff, France).

## Author contributions

ASL, ECM, and AC designed experiments and wrote the manuscript.

ASL and AC cloned, expressed, purified sulfatases and performed the enzymatic assays.

AC, DPB, JAL, and PAE carried out and analysed kinetic and binding experiments EY and JAL performed labelling and NMR experiments

AC and AB performed structural biology experiments. MC and TB performed sulfatase phylogenetic analyses.

GW carried out light scattering experiments and size determination. All authors read and approved the manuscript.

Supplemental Figure 1. Extracted ligands and their electron density maps. The 2*mF*_obs_-*F*_c_ maps are shown contoured at 1σ for all substrates and products co-crystallised with their respective sulfatase.

Supplemental Figure 2. Biophysical and kinetics analysis of BT1918^3S-GlcNAc^ **a**, Right panel shows chromatogram of size exclusion chromatography coupled to light scattering for BT1918^3S-GlcNAc^ demonstrating the presence of a dimer(e= expected molecular mass; o=observed molecular mass). Left panel shows surface representation of the dimer of BT1918^3S-GlcNAc^; the active site is highlighted with a black dashed circle. **b**, Thin layer chromatography analysis of BT1918^3S-GlcNAc^ and its mutants versus its 3S, 6S-*N*-acetylglucosamine substrate. Assays were ran for 48 h, at 37°C, deploying 6 mM substrate and 5 μM enzyme. All assays contained 3 mM HEPES pH 7.0, 45 mM NaCl, and 5 mM CaCl_2_. **c**, DSF analysis showing the stability of the mutant proteins of BT1918^3S-GlcNAc^ with respect to the wildtype (WT). **d)** Thin layer chromatography analysis of BT1918^3S-GlcNAc^ versus 3S-glucosamine and 3S,6S-glucosamine. Assays were ran for 48 h, at 37°C, deploying 6 mM substrate and 5 μM enzyme. All assays contained 3 mM HEPES pH 7.0, 45 mM NaCl, and 5 mM CaCl_2_.

Supplemental Figure 3. Gel chromatography and light scattering of S1_16 sulfatases. Top panels are size exclusion chromatograms whilst, the middle and bottom panels are size exclusion coupled to light scattering chromatograms for the monomer (green) and dimer (black) peaks purified from size exclusion. e= expected molecular mass; o=observed molecular mass.

Supplemental Figure 4. Activity and stability analysis of S1_16 sulfatases and their mutant variants. **a**, DSF analysis of the effects of galactose and *N*-acetyl galactosamine on thermostability with a positive shift indicative of binding. **b**, Normalised DSF melt curves of BT3057^4S-Gal/GalNAc^ and BT3796^4S-Gal/GalNAc^. **c**, DSF melt curves of the purified monomer and dimer species (left), monomer species in the presence of galactose and *N*-acetyl galactosamine (middle), and the dimer species in the presence of galactose and *N*-acetyl galactosamine (right). **d**, Thin layer chromatography (TLC) analysis of wildtype (WT) and mutant S1_16 sulfatases. Asterisks are placed above lanes where activity is observed. **e**, High pressure anion exchange chromatography (HPAEC) of wildtype (WT) and mutant S1_16 sulfatases. A grey block highlights the produced product. Both TLC and HPAEC reactions utilised 6 mM substrate and 1 μM enzyme, except for W109A variants where 10 μM was deployed, with 3 mM HEPES, 45 mM NaCl, and 5 mM CaCl2 over a 48 h period at 37°C.

Supplemental Figure 5. Gel chromatography and light scattering of S1_15 and S1_11 sulfatases. **a**, Chromatograms of size exclusion chromatography coupled to light scattering for S1_15 subfamily members; **b**, Chromatograms of size exclusion chromatography coupled to light scattering for S1_11 subfamily members. e= expected molecular mass; o=observed molecular mass.

Supplemental Figure 6. Analysis of the activity and stability of BT1624^6S-Gal/GalNAc^ and its mutant variants. **a**, Thin layer chromatography (TLC) analysis of wildtype (WT) BT1624^6S-Gal/GalNAc^ and its mutants. Asterisks are placed above lanes where activity is observed. Both TLC reactions utilised 6 mM substrate and 5 μM enzyme, with 3 mM HEPES, 45 mM NaCl, and 5 mM CaCl2 over a 48 h period at 37°C. **b**, DSF analysis showing the stability of the mutant proteins of BT1624^6S-Gal/GalNAc^ with respect to the wildtype. **c**, DSF analysis of the effects of alanine scanning on the ability of BT1624^6S-Gal/GalNAc^ to bind galactose with the T_m_ of the protein shown atop the bar. The experiments were performed with 5 μM of protein and 324 mM of galactose in 100 mM BTP and 150 mM NaCl.

Supplemental Figure 7. **Biochemical comparison of S1_15 BT3333^6S-GalNAc^ and S1_11 sulfatase BT4656^6S-GlcNAc/GlcNS^ a**, Isothermal titration calorimetry traces of inactive serine variants of BT3333^6S-GalNAc^ and BT4656^6S-GlcNAc/GlcNS^ binding to their respective 6S-GalNAc and 6S-GlcNAc substrates. **b**, Specific activity of BT3333^6S-GalNAc^ and BT4656^6S-GlcNAc^ against 0.5 mM their respective 6S-GalNAc and 6S-GlcNAc substrates, monitored by 1D H-NMR. **c**, Acid hydrolysis of 10 mM 6S-GalNAc and 6S-GlcNAc in the presence of 250 mM HCl at 90°C.

Supplemental Figure 8. Full Phylogenetic tree of representative sulfatases from subfamily S1_46. The S1_46 subfamily is composed of 574 sequences (3 eukaryota, 5 Archaea and 566 Bacteria). To avoid identical sequences, and based on taxonomic diversity, 356 sequences were selected for alignment. 15 sequences that belong to the S1_0 subfamily (Phosphonate monoester hydrolase / phosphodiesterase) have been used as outgroup for the alignment.

Supplemental Figure 9. Full Phylogenetic tree of representative sulfatases from subfamily S1_16. The S1_16 subfamily is composed of 1356 sequences. To avoid the identical sequences, and based on taxonomic diversity, 800 representative sequences were used for the alignment. 15 sequences that belong to the S1_0 subfamily (Phosphonate monoester hydrolase / phosphodiesterase) have been used as outgroup for the alignment.

Supplemental Figure 10. Full Phylogenetic tree of representative sulfatases from subfamily S1_15. The S1_15 sub-family is composed of 1895 sequences. To avoid the identical sequences, and based on taxonomic diversity 920 representative sulfatase sequence were selected. 15 sequences that belong to the S1_0 subfamily (Phosphonate monoester hydrolase / phosphodiesterase) have been used as outgroup for the alignment.

Supplemental Figure 11. Full Phylogenetic tree of representative sulfatases from subfamily S1_11. Based on taxonomic diversity, 955 representative sulfatase sequences of the S1_11 subfamily (44% of the subfamily) was aligned. There are 15 sequences of the S1_0 subfamily, phosphonate monoesters, used as outgroup for phylogeny.

---

**Extended data 1.**
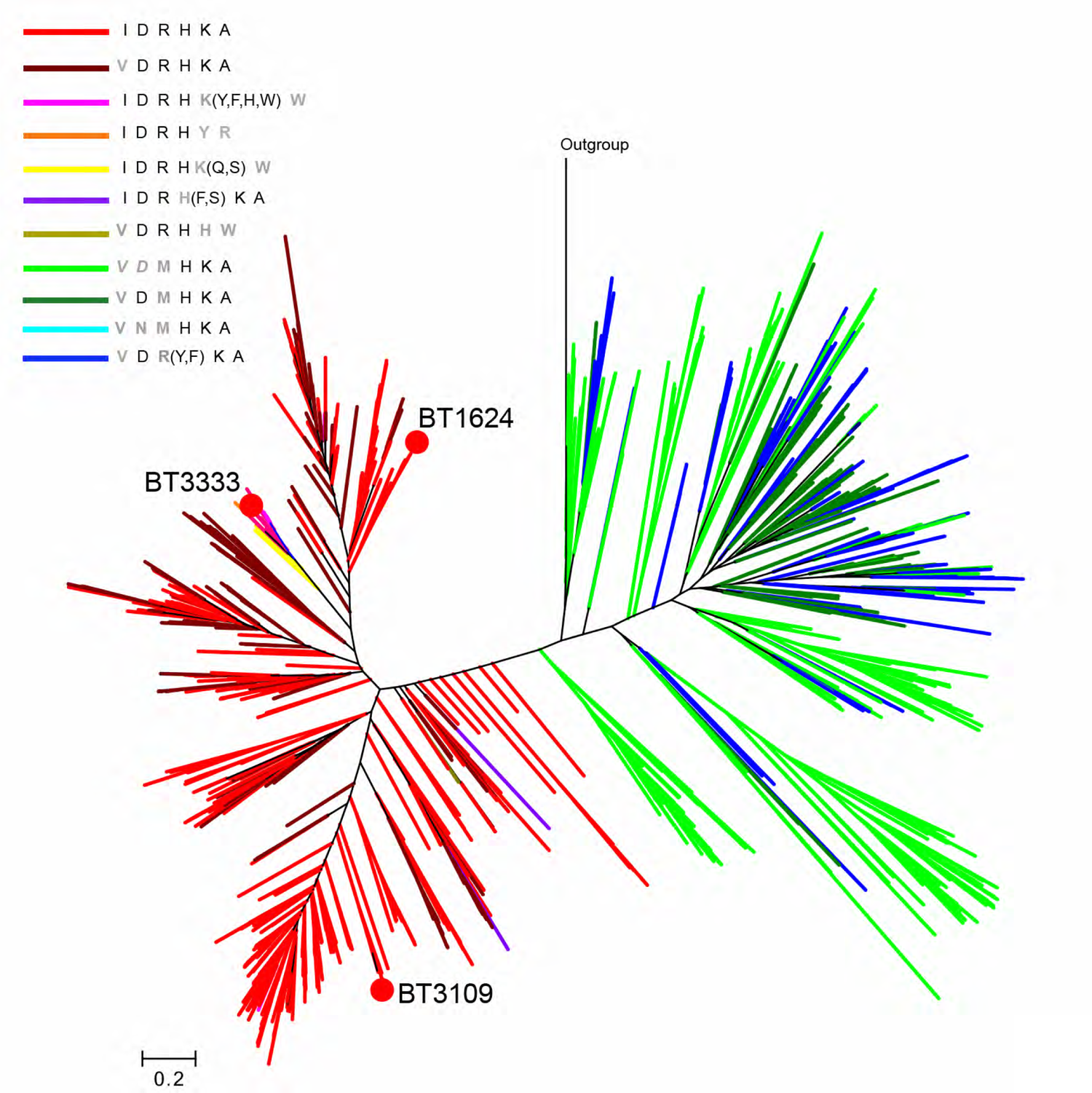
Radial version of the phylogenetic tree of representative sulfatases from subfamily S1_15. For clarity all labels and sequence accession codes have been omitted. The annotations next to the colour code concern the presence or absence of conservation of the indicated residues and in this order: I100, D170, R171, H220, K461 and A462. These residues are crucial in substrate recognition by BT1624^6S-Gal/ GalNAc^ (acc-code Q8A7A1). For simplification the residue numbers have been omitted. For example, an I in black means an equivalent isoleucine is present; a grey and bold letter at any position means that the corresponding residue is replaced by that amino acid; a grey and italic letter at any position means that the equivalent position can be replaced by any type of amino acid; a bold grey letter followed by one-letter codes in parentheses indicates that the equivalent position is substituted by any of those amino acids. Branches having the same colour have the corresponding pattern in common. Red filled circles designate sequences of S1_15 sulfatases from *B. thetaiotaomicron* (See Figure S10 for full tree).

**Extended data 2.**
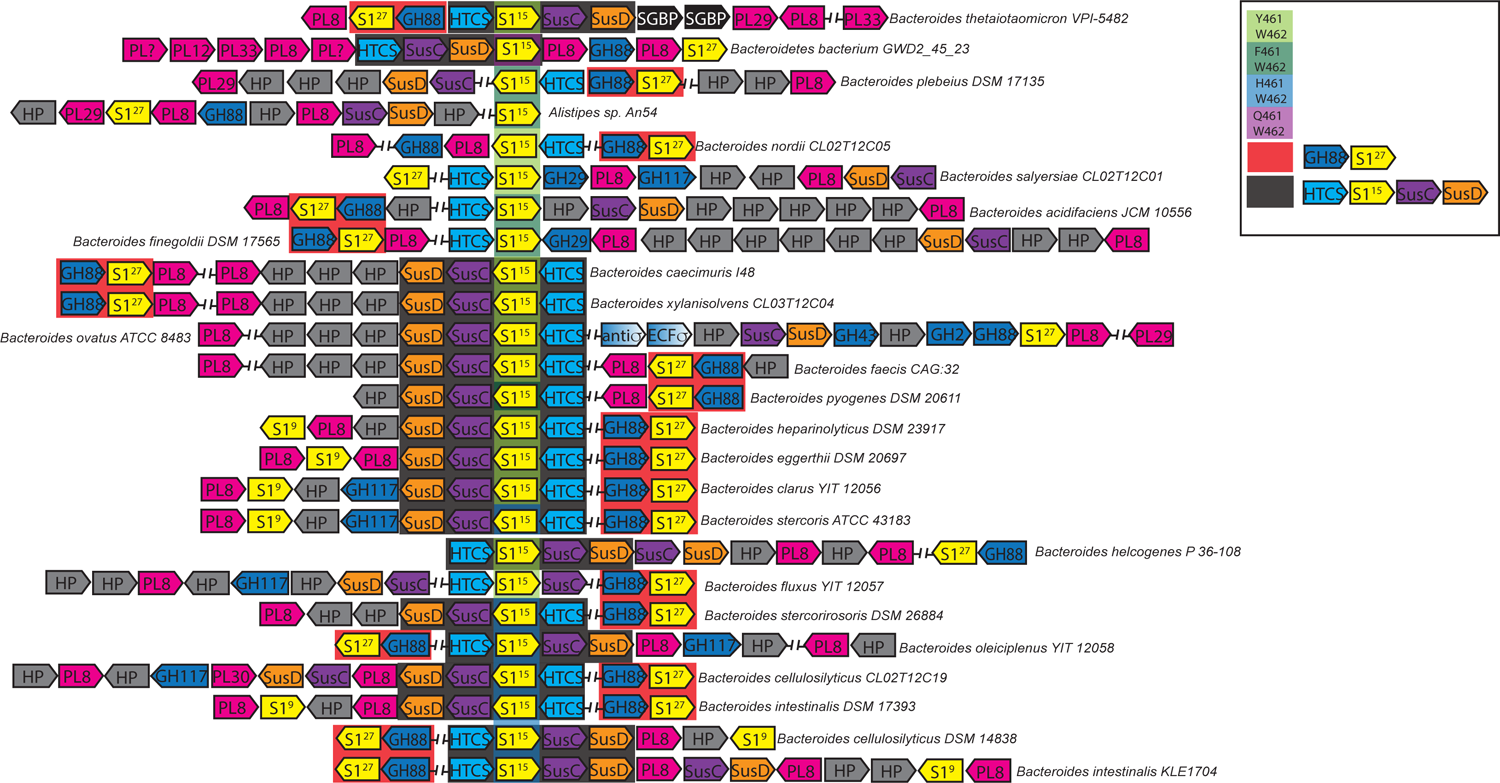
Analysis of S1_15 enzymes with PULs targeting chondroitin sulfate. PULs targeting chondroitin sulfate aligned by orthologues of BT3333^6S-GalNAc^. Light green background shows orthologues with Y463/W464, a dark green background highlights orthologues with F463/W464, a light blue back ground highlight orthologues with H463/W464, and a purple background highlight orthologues with Q463/W464. The numbering used corresponds to the sequence of BT3333^6S-GalNAc^. A red background highlights the presence of GH88 and S1_27 (an endo 4S-chrondroitin sulfatase) which appear to be discrete genetic block not always physically localised to the PUL. A black background highlights a core block observed in CS PULs containing BT3333^6S-GalNAc^ orthologues.

**Extended data 3.**
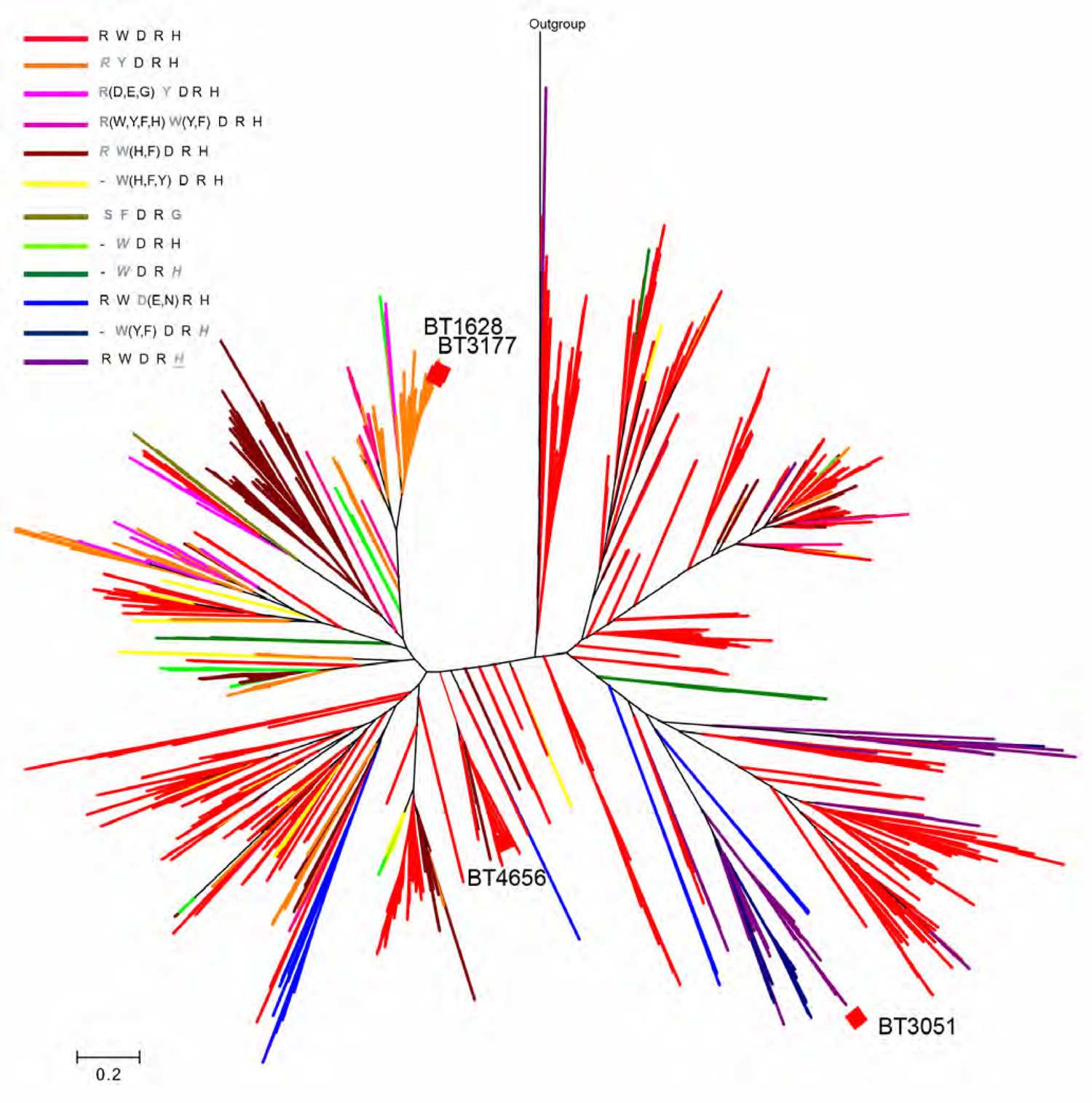
Radial version of the phylogenetic tree of representative sulfatases from subfamily S1_11 For clarity all labels and sequence accession codes have been omitted. The annotations next to the colour code concern the presence or absence of conservation of the indicated residues and in this order: R290, W273, D385, R387 and H471. These residues are crucial in substrate recognition by BT4656^6S-GlcNAc/GlcNS^ (acc-code Q89YS5). For simplification the residue numbers have been omitted. For example, a R in black means an equivalent arginine is present; a grey and bold letter at this position means that the corresponding residue is replaced by that amino acid; the grey and italic R at this position means that the R-equivalent position is replaced by any type of amino acid; a bold grey R followed by one-letter codes in parentheses indicates that the R-equivalent position can be substituted by any of those amino acids; the dash at the R-equivalent position indicates that no equivalent amino acid can be deduced from the multiple alignment. Branches having the same colour have the corresponding pattern in common. Red filled diamonds designate sequences of S1_11 sulfatases from *B. thetaiotaomicron*. All sequences in the specific branch that contains BT4656^6S-GlcNAc/GlcNS^ are found within a conserved heparin PUL (See Figure S11 for full tree).

**Extended data 4.**
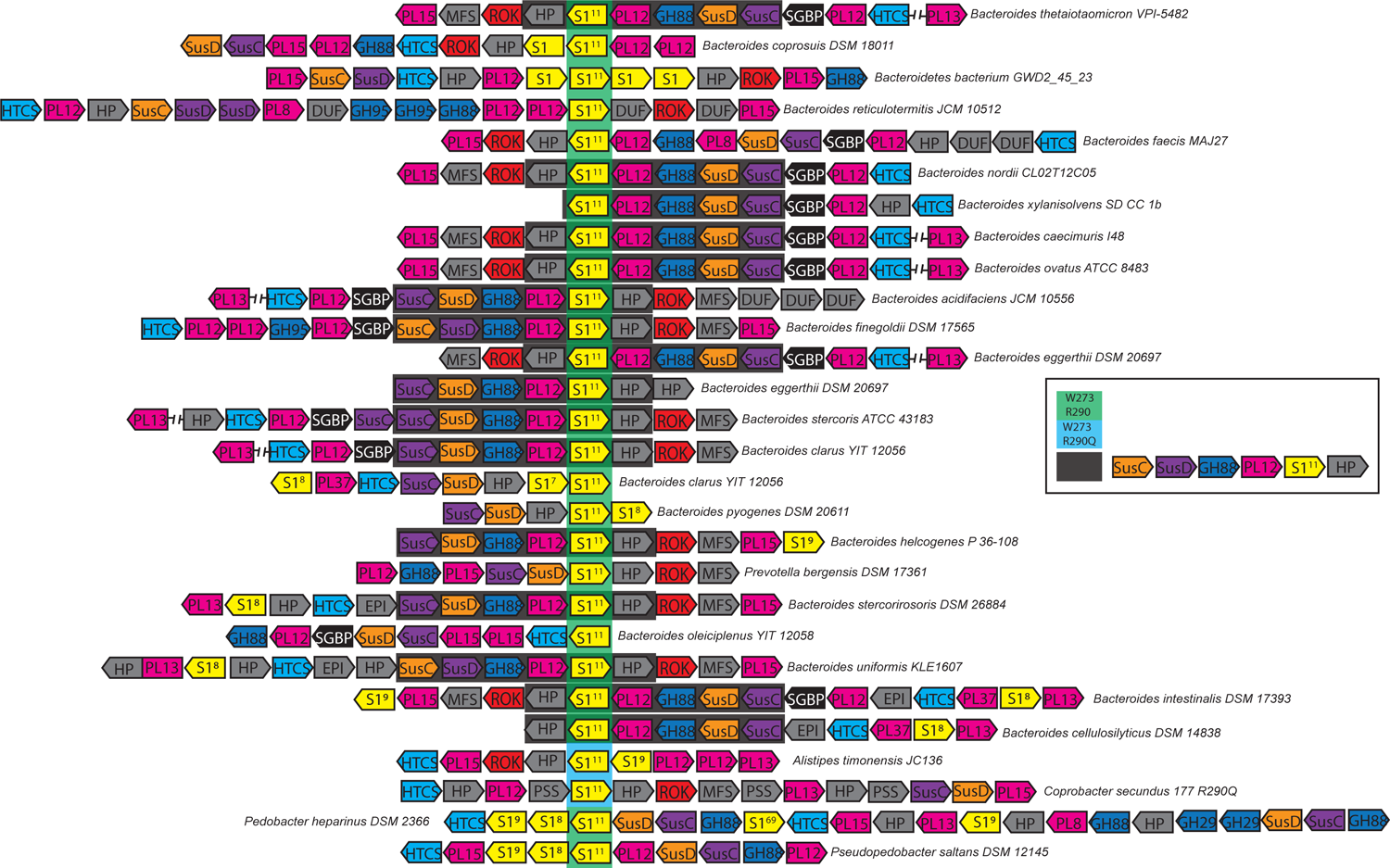
Analysis of S1_11 enzymes with PULs targeting chondroitin sulfate. PULs targeting heparan sulfate (HS) aligned by orthologues of BT4656^6S-GlcNAc/GlcNS^. Orthologues of BT4656^6S-GlcNAc/GlcNS^ with W273/R290 and W273/ Q290 are highlighted with green and blue background, respectively. A black background highlights a core block observed in HS PULs containing BT4656^6S-GlcNAc/GlcNS^ orthologues.

## References

1. Sender, R., Fuchs, S. & Milo, R. Revised Estimates for the Number of Human and Bacteria Cells in the Body. PLoS biology 14, e1002533, doi:10.1371/journal.pbio.1002533 (2016).

2. Cheng, H. Y., Ning, M. X., Chen, D. K. & Ma, W. T. Interactions Between the Gut Microbiota and the Host Innate Immune Response Against Pathogens. Front Immunol 10, 607, doi:10.3389/fimmu.2019.00607 (2019).

3. Belkaid, Y. & Hand, T. W. Role of the microbiota in immunity and inflammation. Cell 157, 121–141, doi:10.1016/j.cell.2014.03.011 (2014).

4. McNeil, N. I. The contribution of the large intestine to energy supplies in man. The American journal of clinical nutrition 39, 338–342, doi:10.1093/ajcn/39.2.338 (1984).

5. Larsbrink, J. et al. A discrete genetic locus confers xyloglucan metabolism in select human gut Bacteroidetes. Nature 506, 498–502, doi:10.1038/nature12907 (2014).

6. Luis, A. S. et al. Dietary pectic glycans are degraded by coordinated enzyme pathways in human colonic Bacteroides. Nature microbiology 3, 210-219, doi:10.1038/s41564-017-0079-1 (2018).

7. Cartmell, A. et al. A surface endogalactanase in Bacteroides thetaiotaomicron confers keystone status for arabinogalactan degradation. Nature microbiology 3, 1314-1326, doi:10.1038/s41564-018-0258-8 (2018).

8. Cartmell, A. et al. How members of the human gut microbiota overcome the sulfation problem posed by glycosaminoglycans. Proceedings of the National Academy of Sciences of the United States of America 114, 7037–7042, doi:10.1073/pnas.1704367114 (2017).

9. Ndeh, D. et al. Metabolism of multiple glycosaminoglycans by Bacteroides thetaiotaomicron is orchestrated by a versatile core genetic locus. Nature communications 11, 646, doi:10.1038/s41467-020-14509-4 (2020).

10. Luis, A. S. et al. A single bacterial sulfatase is required for metabolism of colonic mucin O-glycans and intestinal colonization by a symbiotic human gut bacterium. *bioRxiv*, 2020.2011.2020.392076, doi:10.1101/2020.11.20.392076 (2020).

11. El Kaoutari, A., Armougom, F., Gordon, J. I., Raoult, D. & Henrissat, B. The abundance and variety of carbohydrate-active enzymes in the human gut microbiota. Nat Rev Microbiol 11, 497–504, doi:10.1038/nrmicro3050 (2013).

12. Martens, E. C., Chiang, H. C. & Gordon, J. I. Mucosal glycan foraging enhances fitness and transmission of a saccharolytic human gut bacterial symbiont. Cell host & microbe 4, 447–457, doi:10.1016/j.chom.2008.09.007 (2008).

13. Terrapon, N., Lombard, V., Gilbert, H. J. & Henrissat, B. Automatic prediction of polysaccharide utilization loci in Bacteroidetes species. Bioinformatics 31, 647–655, doi:10.1093/bioinformatics/btu716 (2015).

14. Goodman, A. L. et al. Identifying genetic determinants needed to establish a human gut symbiont in its habitat. Cell host & microbe 6, 279–289, doi:10.1016/j.chom.2009.08.003 (2009).

15. Li, H. et al. The outer mucus layer hosts a distinct intestinal microbial niche. Nature communications 6, 8292, doi:10.1038/ncomms9292 (2015).

16. Tuncil, Y. E. et al. Reciprocal Prioritization to Dietary Glycans by Gut Bacteria in a Competitive Environment Promotes Stable Coexistence. mBio 8, doi:10.1128/mBio.01068-17 (2017).

17. Raghavan, V. & Groisman, E. A. Species-specific dynamic responses of gut bacteria to a mammalian glycan. J Bacteriol 197, 1538–1548, doi:10.1128/JB.00010-15 (2015).

18. Tsai, H. H., Dwarakanath, A. D., Hart, C. A., Milton, J. D. & Rhodes, J. M. Increased faecal mucin sulphatase activity in ulcerative colitis: a potential target for treatment. Gut 36, 570–576, doi:10.1136/gut.36.4.570 (1995).

19. Alipour, M. et al. Mucosal Barrier Depletion and Loss of Bacterial Diversity are Primary Abnormalities in Paediatric Ulcerative Colitis. J Crohns Colitis 10, 462–471, doi:10.1093/ecco-jcc/jjv223 (2016).

20. Kang, S. S. et al. An antibiotic-responsive mouse model of fulminant ulcerative colitis. PLoS Med 5, e41, doi:10.1371/journal.pmed.0050041 (2008).

21. Hickey, C. A. et al. Colitogenic Bacteroides thetaiotaomicron Antigens Access Host Immune Cells in a Sulfatase-Dependent Manner via Outer Membrane Vesicles. Cell host & microbe 17, 672–680, doi:10.1016/j.chom.2015.04.002 (2015).

22. Barbeyron, T. et al. Matching the Diversity of Sulfated Biomolecules: Creation of a Classification Database for Sulfatases Reflecting Their Substrate Specificity. PloS one 11, e0164846, doi:10.1371/journal.pone.0164846 (2016).

23. Hanson, S. R., Best, M. D. & Wong, C. H. Sulfatases: structure, mechanism, biological activity, inhibition, and synthetic utility. Angewandte Chemie 43, 5736–5763, doi:10.1002/anie.200300632 (2004).

24. Hettle, A. G. et al. The Molecular Basis of Polysaccharide Sulfatase Activity and a Nomenclature for Catalytic Subsites in this Class of Enzyme. Structure 26, 747–758 e744, doi:10.1016/j.str.2018.03.012 (2018).

25. Lapebie, P., Lombard, V., Drula, E., Terrapon, N. & Henrissat, B. Bacteroidetes use thousands of enzyme combinations to break down glycans. Nature communications 10, 2043, doi:10.1038/s41467-019-10068-5 (2019).

26. Pudlo, N. A. et al. Extensive transfer of genes for edible seaweed digestion from marine to human gut bacteria. *bioRxiv*, 2020.2006.2009.142968, doi:10.1101/2020.06.09.142968 (2020).

27. Roche, P. et al. Molecular basis of symbiotic host specificity in Rhizobium meliloti: nodH and nodPQ genes encode the sulfation of lipo-oligosaccharide signals. Cell 67, 1131–1143, doi:10.1016/0092-8674(91)90290-f (1991).

28. Byrne, D. P. et al. cAMP-dependent protein kinase (PKA) complexes probed by complementary differential scanning fluorimetry and ion mobility-mass spectrometry. The Biochemical journal 473, 3159–3175, doi:10.1042/BCJ20160648 (2016).

29. Das, T. M., Rao, C. P. & Kolehmainen, E. Synthesis and characterisation of N-glycosyl amines from the reaction between 4,6-O-benzylidene-D-glucopyranose and substituted aromatic amines and also between 2-(o-aminophenyl)benzimidazole and pentoses or hexoses. Carbohydr Res 334, 261–269, doi:10.1016/s0008-6215(01)00202-6 (2001).

30. Kabsch, W. Xds. Acta crystallographica. Section D, Biological crystallography 66, 125–132, doi:10.1107/S0907444909047337 (2010).

31. Evans, P. Scaling and assessment of data quality. Acta crystallographica. Section D, Biological crystallography 62, 72–82, doi:10.1107/S0907444905036693 (2006).

32. Evans, P. R. An introduction to data reduction: space-group determination, scaling and intensity statistics. Acta crystallographica. Section D, Biological crystallography 67, 282–292, doi:10.1107/S090744491003982X (2011).

33. Long, F., Vagin, A. A., Young, P. & Murshudov, G. N. BALBES: a molecular-replacement pipeline. Acta crystallographica. Section D, Biological crystallography 64, 125–132, doi:10.1107/S0907444907050172 (2008).

34. McCoy, A. J. Solving structures of protein complexes by molecular replacement with Phaser. Acta crystallographica. Section D, Biological crystallography 63, 32–41, doi:10.1107/S0907444906045975 (2007).

35. Emsley, P., Lohkamp, B., Scott, W. G. & Cowtan, K. Features and development of Coot. Acta Crystallogr D Biol Crystallogr 66, 486–501, doi:10.1107/S0907444910007493 (2010).

36. Murshudov, G. N. et al. REFMAC5 for the refinement of macromolecular crystal structures. Acta crystallographica. Section D, Biological crystallography 67, 355–367, doi:10.1107/S0907444911001314 (2011).

37. Lebedev, A. A. et al. JLigand: a graphical tool for the CCP4 template-restraint library. Acta crystallographica. Section D, Biological crystallography 68, 431–440, doi:10.1107/S090744491200251X (2012).

38. Chen, V. B. et al. MolProbity: all-atom structure validation for macromolecular crystallography. *Acta crystallographica. Section D*, Biological crystallography 66, 12–21, doi:10.1107/S0907444909042073 (2010).

39. Potterton, L. et al. CCP4i2: the new graphical user interface to the CCP4 program suite. Acta Crystallogr D Struct Biol 74, 68–84, doi:10.1107/S2059798317016035 (2018).

40. Collaborative Computational Project, N. The CCP4 suite: programs for protein crystallography. Acta crystallographica. Section D, Biological crystallography 50, 760–763, doi:10.1107/S0907444994003112 (1994).

41. Katoh, K., Misawa, K., Kuma, K. & Miyata, T. MAFFT: a novel method for rapid multiple sequence alignment based on fast Fourier transform. Nucleic Acids Res 30, 3059–3066, doi:10.1093/nar/gkf436 (2002).

42. Clamp, M., Cuff, J., Searle, S. M. & Barton, G. J. The Jalview Java alignment editor. Bioinformatics 20, 426–427, doi:10.1093/bioinformatics/btg430 (2004).

43. Stamatakis, A. RAxML version 8: a tool for phylogenetic analysis and post-analysis of large phylogenies. Bioinformatics 30, 1312–1313, doi:10.1093/bioinformatics/btu033 (2014).

44. Felsenstein, J. Evolutionary trees from DNA sequences: a maximum likelihood approach. J Mol Evol 17, 368–376, doi:10.1007/BF01734359 (1981).

45. Le, S. Q. & Gascuel, O. An improved general amino acid replacement matrix. Mol Biol Evol 25, 1307–1320, doi:10.1093/molbev/msn067 (2008).

46. Felsenstein, J. Confidence Limits on Phylogenies: An Approach Using the Bootstrap. Evolution 39, 783–791, doi:10.1111/j.1558-5646.1985.tb00420.x (1985).

47. Varki, A. et al. Symbol Nomenclature for Graphical Representations of Glycans. Glycobiology 25, 1323–1324, doi:10.1093/glycob/cwv091 (2015).

