## Supplemental discussion and figures for "Sulfated host glycan recognition by carbohydrate sulfatases of the human gut microbiota"

*Correspondence to:

### Supplemental discussion

*Comparison of S1_11 and S1_15 biochemical properties*

The three residues (Asp, Arg and His) that form a 'recognition triad' in the S1_11 sulfatases to recognise the O3 and O4 of GlcNAc are the same as those used by S1_15 members to coordinate the O3 and O4 of GalNAc, Therefore, both subfamilies appear to utilise a comparable binding mechanism. Indeed, inactive forms of BT4656$^{6S\text{-}GlcNAc/GlcNS}$ and BT3333$^{6S\text{-}GalNAc}$ have similar affinities, and binding energy, for their respective substrates, (**Figure S7a and Table S8**). Yet, the enzymes display vastly different catalytic rates, with the S1_15 enzyme BT3333$^{6S\text{-}GalNAc}$ displaying a specific activity >100 lower than the S1_11 subfamily member BT4656$^{6S\text{-}GlcNAc/GlcNS}$ (**Figure S7b**). In addition, there was no significant difference in the reactivity of the their 6S-GlcNAc and 6S-GalNAc substrates under acid hydrolysis conditions (**Figure S7c** and **Table S8**), suggesting that the difference in rate may be due to the ability of the *E.coli* expression host to correctly insert the catalytic FGly residue into S1_15 subfamily members at levels comparable to other subfamilies characterised. Interestingly although BT3333$^{6S\text{-}GalNAc}$ and BT4656$^{6S\text{-}GlcNAc/GlcNS}$ bound their respective *N*-acetylated sugar with similar affinities BT3333$^{6S\text{-}GalNAc}$ displayed a significant entropic contribution to binding whilst, BT4656$^{6S\text{-}GlcNAc/GlcNS}$ was entirely enthalpy driven (**Table S8**). These differences likely reflect the variation with how the two enzymes interact with the C2 substituent of their substrates with BT3333$^{6S\text{-}GalNAc}$ utilising a hydrophobic interaction whilst BT4656$^{6S\text{-}GlcNAc/GlcNS}$ utilises an ionic interaction. This further demonstrates how the non-conserved regions of S1 subfamilies effect not just the substrate specificity but the overall thermodynamics of the interaction too.

**Table S1**

Catalytic activities of S1_46 and S1_16 enzymes and their mutant variants

| Sulfatase S1 subfamily | Enzyme/substrate | 3S,6S-GlcNAc | 4S-Gal | 4S-GalNac |
|---|---|---|---|---|
| S1_46 | BT1918$^{3S\text{-}Gal}$ S73C (WT) | (6.1 ± 0.4) x $10^4$ | NA | NA |
| | BT1918$^{3S\text{-}Gal}$ Y94A | (1.5 ± 0.02) x $10^3$ | NT | NT |
| | BT1918$^{3S\text{-}Gal}$ Y94F | (3.1 ± 0.3) x $10^4$ | NT | NT |
| | BT1918$^{3S\text{-}Gal}$ R143A | (1.4 ± 0.1) x $10^3$ | NT | NT |
| | BT1918$^{3S\text{-}Gal}$ W173A | (4.5 ± 0.4) x $10^4$ | NT | NT |
| | BT1918$^{3S\text{-}Gal}$ N174A | (2.3 ± 0.1) x $10^3$ | NT | NT |
| | BT1918$^{3S\text{-}Gal}$ R327A | (6.8 ± 0.6) x $10^3$ | NT | NT |
| | BT1918$^{3S\text{-}Gal}$ Y408A | (3.1 ± 0.6) x $10^3$ | NT | NT |
| | BT1918$^{3S\text{-}Gal}$ Y408F | (4.7 ± 0.2) x $10^4$ | NT | NT |
| S1_16 | BT3057$^{4S\text{-}Gal/GalNAc}$ S88C (WT) | NA | (4.2 ± 0.4) x $10^3$ | (5.0 ± 0.1) x $10^3$ |
| | BT3057$^{4S\text{-}Gal/GalNAc}$ W109A | NA | NQ | NQ |
| | BT3057$^{4S\text{-}Gal/GalNAc}$ H182A | NT | (3.4 ± 0.3) x $10^2$ | (4.4 ± 0.3) x $10^2$ |
| | BT3057$^{4S\text{-}Gal/GalNAc}$ H423A | NT | (2.6 ± 0.2) x $10^2$ | (2.4 ± 0.5) x $10^2$ |
| S1_16 | BT3796$^{4S\text{-}Gal/GalNAc}$ S88C (WT) | NA | (2.7 ± 0.2) x $10^4$ | (2.2 ± 0.4) x $10^4$ |
| | BT3796$^{4S\text{-}Gal/GalNAc}$ W109A | NT | NQ | NQ |
| | BT3796$^{4S\text{-}Gal/GalNAc}$ W332A | NT | (1.0 ± 0.1) x $10^3$ | (3.3 ± 0.1) x $10^2$ |
| | BT3796$^{4S\text{-}Gal/GalNAc}$ W431A | NT | (1.3 ± 0.1) x $10^3$ | (1.8 ± 0.4) x $10^2$ |

Data represent technical triplicates and the errors are reported as standard errors of the mean. Assays were performed in Bis-Tris propane at pH 7, 6.5, and 8.5 for BT1918$^{3S\text{-}Gal}$, BT3057$^{4S\text{-}Gal/GalNAc}$, and BT3796$^{4S\text{-}Gal/GalNAc}$, respectively, and where supplemented with 150 mM NaCl, 5 mM CaCl2, and 0.02 % (v/v) Brij-35. The pH optima of the enzymes was established in previous work[1]. WT, wildtype enzymes; NA, not active; NT, not tested; NQ, activity was not quantifiable.

### Table S2

**S1_46 sequences detailing protein accession number, species, phylum, and the environmental context of each sequence.**
Sequences are arranged in the same order as the phylogenetic tree (Figure S8). This table is provided separately as excel spreadsheet with multitabs for the different bacteria environments.

### Table S3

**S1_16 sequences detailing protein accession number, species, phylum, and the environmental context of each sequence.**
Sequences are arranged in the inverse order as the phylogenetic tree (Figure S9). That is the bottom of the phylogenetic tree in Figure S9 corresponds to the top of the excel spreadsheet. This table is provided separately as excel spreadsheet with multitabs for the different bacteria environments.

### Table S4

**S1_15 sequences detailing protein accession number, species, phylum, and the environmental context of each sequence.**
Sequences are arranged in the same order as the phylogenetic tree (Figure S10). This table is provided separately as excel spreadsheet with multitabs for the different bacteria environments.

**Table S5**

Catalytic activities of S1_15 and S1_11 enzymes and their mutant variants

| Sulfatase S1 subfamily | Enzyme/substrate | 6S-Gal* | 6S-GalNAc* | 6S-GlcNAc | 6S-GlcNS |
|---|---|---|---|---|---|
| **S1_15** | BT3109$^{6S-Gal/GalNAc}$ S73C (WT) | 2.73 ± 0.74 | NQ | NA | NA |
| **S1_15** | BT3333$^{6S-GalNAc}$ S80C (WT) | NQ | 3.1 ± 0.02 | NA | NA |
| **S1_15** | BT4631$^{6S-Gal/GalNAc}$ S85C (WT) | NQ | NQ | NA | NA |
| **S1_15** | BT1624$^{6S-Gal/GalNAc}$ S79C (WT) | 3.9 ± 0.55 | 1.95 ± 0.6 | NA | NA |
| | BT1624$^{6S-Gal/GalNAc}$ D170A | NQ | NQ | NT | NT |
| | BT1624$^{6S-Gal/GalNAc}$ R171A | NQ | NQ | NT | NT |
| | BT1624$^{6S-Gal/GalNAc}$ H220A | NQ | NQ | NT | NT |
| | BT1624$^{6S-Gal/GalNAc}$ D336A | NQ | NQ | NT | NT |
| | BT1624$^{6S-Gal/GalNAc}$ Y360A | NQ | NQ | NT | NT |
| | BT1624$^{6S-Gal/GalNAc}$ Y360F | NQ | NQ | NT | NT |
| **S1_11** | BT4656$^{6S-GlcNAc/GlcNS}$ S91C (WT) | | | $(5.4 \pm 0.3) \times 10^4$ | $(6.4 \pm 0.6) \times 10^4$ |
| **S1_11** | BT3177$^{6S-GlcNAc}$ S79C (WT) | NA | NA | $(6.3 \pm 0.2) \times 10^3$<br>$(2.6 \pm 0.2) \times 10^4$ | $(6.1 \pm 0.6) \times 10^3$ |
| | BT3177$^{6S-GlcNAc}$ Q147A | NT | NT | $(5.9 \pm 0.2) \times 10^3$ | NT |
| | BT3177$^{6S-GlcNAc}$ R202A | NA | NA | NA | NT |
| | BT3177$^{6S-GlcNAc}$ Y250A | NT | NT | $(5.4 \pm 0.2) \times 10^2$ | NT |
| | BT3177$^{6S-GlcNAc}$ Y250F | NT | NT | $(7.8 \pm 0.6) \times 10^3$ | NT |
| | BT3177$^{6S-GlcNAc}$ D361A | NT | NT | NA | NT |
| | BT3177$^{6S-GlcNAc}$ R363A | NT | NT | $(1.8 \pm 0.1) \times 10^3$ | NT |
| | BT3177$^{6S-GlcNAc}$ E334A | NT | NT | $(5.6 \pm 2.2) \times 10^0$ | NT |
| | BT3177$^{6S-GlcNAc}$ H445A | NT | NT | $(1.8 \pm 0.2) \times 10^1$ | NT |

Data represent technical triplicates and the errors are reported as standard errors of the mean. Assays for BT3177$^{6S-GlcNAc}$ was performed in Bis-Tris propane at pH 8, and MES at pH 6.0 for BT4656$^{6S-GlcNAc/GlcNS}$ and where supplemented with 150 mM NaCl, 5 mM CaCl2, and 0.02 % (v/v) Brij-35. The pH optima of the S1_11 enzymes was established in the previous works[1]. *indicates that these are specific activities and the data are reproduced from a previous publication[1]. For BT3177$^{6S-GlcNAc}$ blue values were carried out at the same as time assays for BT4656$^{6S-GlcNAc/GlcNS}$ whilst black values represent mutants and wildtype experiments conducted at the same time. WT, wildtype; NA, not active; NT, not tested; NQ, activity was not quantifiable.

**Table S6**

Environmental niche of the S1_15 protein sequences bearing the C-terminal motif VEEEPLK.

| Protein accession no. | Species/strain | Phylum | Environment |
|---|---|---|---|
| A0A1I1A1I8_9BACT | Algoriphagus aquimarinus | Bacteroidetes | Marine |
| A0A2W7RKQ3_9BACT | Algoriphagus ratkowskyi | Bacteroidetes | Marine |
| A0A2W7SCF5_9BACT | Algoriphagus chordae | Bacteroidetes | Marine |
| A0A3E0DVL9_9BACT | Algoriphagus antarcticus | Bacteroidetes | Marine |
| A0A1I7DQY9_9BACT | Algoriphagus locisalis | Bacteroidetes | Marine |
| A0A327NZ03_9BACT | Algoriphagus yeomjeoni | Bacteroidetes | Marine |
| L0G416_ECHVK | Echinicola vietnamensis (strain DSM 17526 / LMG 23754 / KMM 6221) | Bacteroidetes | Marine |
| A0A2Z4IDG3_9BACT | Echinicola strongylocentroti | Bacteroidetes | Marine |
| A0A1M5CXJ3_9FLAO | Arenibacter palladensis | Bacteroidetes | Marine |
| A0A1X7KJF1_9FLAO | Arenibacter troitsensis | Bacteroidetes | Marine |
| A0A221UVR5_9FLAO | Arenibacter algicola | Bacteroidetes | Marine |
| A0A327R3X5_9FLAO | Arenibacter echinorum | Bacteroidetes | Marine |
| A0A1G9J4E9_9FLAO | Kriegella aquimaris | Bacteroidetes | Marine |
| A0A1M6LF72_9FLAO | Pseudozobellia thermophila | Bacteroidetes | Marine |
| G0L7K9_ZOBGA | Zobellia galactanivorans (strain DSM 12802 / CCUG 47099 / CIP 106680 / NCIMB 13871 / Dsij) | Bacteroidetes | Marine |
| A0A1N7K1D5_9FLAO | Zobellia uliginosa | Bacteroidetes | Marine |
| A0A3D9L5M9_9BACT | Marinoscillum furvescens DSM 4134 | Bacteroidetes | Marine (Brown sand) |
| A0A0D8JAD3_9BACT | Draconibacterium sediminis | Bacteroidetes | Marine (River) |
| A0A0D8JFM7_9BACT | Draconibacterium sediminis | Bacteroidetes | Marine (River) |
| X5DDW6_9BACT | Draconibacterium orientale | Bacteroidetes | Marine |
| A0A0D8JFL1_9BACT | Draconibacterium sediminis | Bacteroidetes | Marine |
| X5DUJ5_9BACT | Maribacter sp. (strain HTCC2170 / KCCM 42371) | Bacteroidetes | Marine |
| A0A0L8VD50_9BACT | Sunxiuqinia dokdonensis | Bacteroidetes | Marine |
| A0A0L8V2I0_9BACT | Sunxiuqinia dokdonensis | Bacteroidetes | Marine |
| A0A2V4A3S1_9BACT | Marinifilum breve | Bacteroidetes | Marine |
| A0A2V4A373_9BACT | Marinifilum breve | Bacteroidetes | Marine |
| A0A285MWN0_9FLAO | Flagellimonas pacifica | Bacteroidetes | Marine |
| A0A3D9EZ57_9FLAO | Muricauda sp. ARW7G5W | Bacteroidetes | Marine |
| A4AMR0_MARSH | Maribacter sp. (strain HTCC2170 / KCCM 42371) | Bacteroidetes | Marine |
| A0A1G7FAH6_9FLAO | Pricia antarctica | Bacteroidetes | Marine |
| A0A3B0C6L6_9FLAO | Ulvibacterium marinum CCMM003 | Bacteroidetes | Marine |
| A0A1G6YXK5_9FLAO | Pricia antarctica | Bacteroidetes | Marine |
| A0A2D0NC76_9BACT | Lewinella nigricans DSM 23189 = NBRC 102662 | Bacteroidetes | Marine (brown beach sand) |
| A0A2U2B614_9BACT | Marinilabilia sp. WTE | Bacteroidetes | Marine |
| A0A2U2B771_9BACT | Marinilabilia sp. WTE | Bacteroidetes | Marine |
| S7X5W7_9BACT | Cyclobacterium qasimii M12-11B | Bacteroidetes | Marine |
| G0IYU3_CYCMS | Cyclobacterium marinum (strain ATCC 25205 / DSM 745) | Bacteroidetes | Marine microbiome (Coelom of sand dollar) |
| A0A0H4PEX4_9BACT | Cyclobacterium amurskyense | Bacteroidetes | Marine |
| A0A1M7PFZ6_9BACT | Cyclobacterium lianum | Bacteroidetes | Marine |
| A0A1H6T1X9_9BACT | Cyclobacterium halophilum | Bacteroidetes | Marine |
| A0A2Z4IEM5_9BACT | Echinicola strongylocentroti | Bacteroidetes | Marine |
| A0A2G2JTY4_9FLAOO | Leeuwenhoekiella sp. | Bacteroidetes | Marine |
| A0A2G1VPX9_9FLAOO | Leeuwenhoekiella nanhaiensis | Bacteroidetes | Marine |
| A0A1H7SK37_9FLAOO | Maribacter orientalis | Bacteroidetes | Marine |
| A0A1T5BFH0_9FLAOO | Maribacter arcticus | Bacteroidetes | Marine |
| E6X915_CELAD | Cellulophaga algicola (strain DSM 14237 / IC166 / ACAM 630) | Bacteroidetes | Marine |
| A0A0A7K6C8_9FLAO | Cellulophaga baltica 18 | Bacteroidetes | Marine |
| A0A1G7JIH3_9FLAO | Cellulophaga baltica | Bacteroidetes | Marine |
| A0A1M6MQ16_9FLAO | Maribacter aquivivus | Bacteroidetes | Marine |

| | | | |
|---|---|---|---|
| T2KN04_FORAG | Formosa agariphila (strain DSM 15362 / KCTC 12365 / LMG 23005 / KMM 3901) | Bacteroidetes | Marine |
| I3CAX1_9FLAOO | Joostella marina DSM 19592 | Bacteroidetes | Marine |
| F0R9D9_CELLC | Cellulophaga lytica (strain ATCC 23178 / DSM 7489 / JCM 8516 / NBRC 14961 / NCIMB 1423 / VKM B-1433 / Cy l20) | Bacteroidetes | Marine |
| A0A1K1R267_9FLAO | Cellulophaga fucicola | Bacteroidetes | Marine |
| A0A238V9J4_9FLAO | Lutibacter agarilyticus | Bacteroidetes | Marine |
| A0A3G2L6S7_9FLAO | Euzebyella marina | Bacteroidetes | Marine |
| A0A2U2B7T4_9BACT | Marinilabilia sp. WTE | Bacteroidetes | Marine |
| A0A0N8KGA9_9BACT | Algoriphagus marincola HL-49 | Bacteroidetes | Marine |
| A0A142ESJ7_9BACT | Algoriphagus sp. M8-2 | Bacteroidetes | Marine |
| A0A1G5WVB9_9BACT | Algoriphagus alkaliphilus | Bacteroidetes | Marine |
| A0A1G6SNN1_9BACT | Algoriphagus faecimaris | Bacteroidetes | Marine |
| A0A1I2NHE7_9BACT | Algoriphagus hitonicola | Bacteroidetes | Marine |
| A0A1I5IK52_9BACT | Algoriphagus ornithinivorans | Bacteroidetes | Marine |
| A0A327PLD7_9BACT | Algoriphagus yeomjeoni | Bacteroidetes | Marine |
| A0A0K8R1W6_9BACT | Bacteroidales bacterium 6E | Bacteroidetes | Paddy field |
| A0A1M5CI20_9BACT | Mariniphaga anaerophila | Bacteroidetes | Marine |
| A0A399CVA0_9BACT | Mariniphaga sediminis | Bacteroidetes | Marine |
| A0A2E9G6Y9_9RHOB | Thalassobius sp. | Bacteroidetes | Marine |
| R7ZL40_9BACT | Lunatimonas lonarensis | Bacteroidetes | Marine |
| A0A1E5T8J6_9FLAO | Algibacter aquaticus | Bacteroidetes | Marine |
| A0A1B1Y625_9FLAO | Wenyingzhuangia fucanilytica | Bacteroidetes | Marine |
| Q93P98_9BACT | Microscilla sp. PRE1 | Bacteroidetes | - |
| T2KLX1_FORAG | Formosa agariphila (strain DSM 15362 / KCTC 12365 / LMG 23005 / KMM 3901) | Bacteroidetes | Marine |
| A0A1B1Y2V9_9FLAO | Wenyingzhuangia fucanilytica | Bacteroidetes | Marine |
| A0A2S1LIQ8_9FLAO | Flavobacterium faecale | Bacteroidetes | Microbiome (Antarctic penguin) |
| A0A1M7JJP9_9FLAO | Flavobacterium saccharophilum | Bacteroidetes | Marine |
| A0A1I2KLW2_9FLAO | Salegentibacter agarivorans | Bacteroidetes | Marine |
| A0A1Y1CFJ4_9BACT | Labilibaculum antarcticum | Bacteroidetes | Marine |
| A0A2N3HTH2_9BACT | Labilibaculum filiforme | Bacteroidetes | Marine |
| A0A1Y1CFY5_9BACT | Labilibaculum antarcticum | Bacteroidetes | Marine |
| W7YL88_9BACT | Saccharicrinis fermentans DSM 9555 = JCM 21142 | Bacteroidetes | Marine |
| A0A1H4UBX0_9FLAO | Maribacter dokdonensis | Bacteroidetes | Marine |
| A0A1M6GJR7_9FLAO | Pseudozobellia thermophila | Bacteroidetes | Marine |
| A0A1H5SGQ7_9BACT | Algoriphagus boritolerans DSM 17298 = JCM 18970 | Bacteroidetes | Soil (boron rich) |
| A0A1W2H561_9BACT | Aquiflexum balticum DSM 16537 | Bacteroidetes | Marine |
| A0A1N7LRG9_9BACT | Belliella pelovolcani | Bacteroidetes | Soil (Volcano mud) |
| A0A239C1A3_9BACT | Belliella buryatensis | Bacteroidetes | Marine (Lake) |
| A0A2P8ECI8_9BACT | Cecembia rubra | Bacteroidetes | Hot spring |
| A0A3N4GU18_9BACT | Cyclobacteriaceae bacterium YHN15 | Bacteroidetes | - |
| A0A1N7LXK0_9BACT | Belliella pelovolcani | Bacteroidetes | Soil (Volcano mud) |
| A0A239ETG6_9BACT | Belliella buryatensis | Bacteroidetes | Marine (Lake) |
| A0A2G0CAT8_9BACT | Lewinella marina | Bacteroidetes | Marine |
| A0A2D8ARS2_9EURY | Euryarchaeota archaeon | Euryarchaeota (Archea) | Environmental (Ground water) |
| A0A1M6UHM9_9BACT | Reichenbachiella agariperforans | Bacteroidetes | Marine |
| A0A2E2SLQ9_9BACT | Cytophagaceae bacterium | Bacteroidetes | - |
| A0A150AAE2_9BACT | Flammeovirga sp. SJP92 | Bacteroidetes | - |
| A0A1S1YTS2_9BACT | Flammeovirga pacifica | Bacteroidetes | Marine |
| A0A1B1FZL8_9BACT | Flammeovirga sp. MY04 | Bacteroidetes | - |
| A0A2S1LHS8_9FLAO | Flavobacterium faecale | Bacteroidetes | Microbiome (Antarctic penguin) |
| A0A1E4A8T8_9BACT | Cytophagaceae bacterium SCN 52-12 | Bacteroidetes | |
| C6VTS4_DYAFD | Dyadobacter fermentans (strain ATCC 700827 / DSM 18053 / NS114) | Bacteroidetes | Environmental (Maize stems) |

| | | | |
|---|---|---|---|
| A0A1I3D2I5_9SPHI | Parapedobacter indicus | Bacteroidetes | Soil (Hexachlorocyclohexane) |
| A0A362WZT5_9FLAO | Jejuia pallidilutea | Bacteroidetes | Marine |
| A0A150AI20_9BACT | Flammeovirga sp. SJP92 | Bacteroidetes | - |
| A0A1B1FZG5_9BACT | Flammeovirga sp. MY04 | Bacteroidetes | Marine |
| A0A150A9M2_9BACT | Flammeovirga sp. SJP92 | Bacteroidetes | - |
| A0A1S1YUB5_9BACT | Flammeovirga pacifica | Bacteroidetes | Marine |
| A0A1B1G299_9BACT | Flammeovirga sp. MY04 | Bacteroidetes | Marine |
| Q8A346_BACTN | Bacteroides thetaiotaomicron (strain ATCC 29148 / DSM 2079 / NCTC 10582 / E50 / VPI-5482) | Bacteroidetes | Microbiome (Human) |
| A7LY94_BACO1 | Bacteroides ovatus (strain ATCC 8483 / DSM 1896 / JCM 5824 / NCTC 11153) | Bacteroidetes | Microbiome (Human) |
| I9SNI8_BACOV | Bacteroides ovatus CL02T12C04 | Bacteroidetes | Microbiome (Human) |
| A0A1C7H3U9_9BACE | Bacteroides caecimuris | Bacteroidetes | Microbiome (Mouse) |
| I9AEF4_9BACE | Bacteroides xylanisolvens CL03T12C04 | Bacteroidetes | Microbiome (Human) |
| W0EQX0_9BACT | Barnesiella viscericola DSM 18177 | Bacteroidetes | Microbiome (Chicken) |

### Table S7

**S1_11 sequences detailing protein accession number, species, phylum, and the environmental context of each sequence.**

Sequences are arranged in the same order as the phylogenetic tree (Figure S11). This table is provided separately as excel spreadsheet with multitabs for the different bacteria environments.

**Table S8**

Comparative table of the binding energies, specific rates, and substrate reactivity for BT3333$^{6S\text{-}GalNAc}$ and BT4656$^{6S\text{-}GlcNAc/GlcNS}$

| ITC analysis of BT3333$^{S1_15}$ and BT4656$^{S1_11}$ | | | | | | | |
|---|---|---|---|---|---|---|---|
| **S1 subfamily** | **Protein** | **Substrate** | ***K*A (M$^{-1}$)** | **ΔG (kcal mol$^{-1}$)** | **ΔH (kcal mol$^{-1}$)** | **TΔS (kcal mol$^{-1}$)** | ***n*** |
| **S1_15** | BT3333$^{6S\text{-}GalNAc}$ | **6S-GalNAc** | (7.5 ± 0.7) x $10^4$ | -6.7 x $10^3$ | (-2.9 ± 0.9) x $10^3$ | 3.8 x $10^3$ | 0.88 ± 0.02 |
| **S1_11** | BT4656$^{6S\text{-}GlcNAc/GlcNS}$ | **6S-GlcNAc** | (3.7 ± 0.7) x $10^5$ | -7.6 x $10^3$ | (-1.7 ± 0.7) x $10^4$ | -9.4 x $10^3$ | 0.82 ± 0.02 |

| Comparison of acid hydrolysis and enzymatic hydrolysis of GalNAc6S and GlcNAc6S | | | | | |
|---|---|---|---|---|---|
| **Substrate** | **Acid hydrolysis with HCl (integral/h)** | **Acid hydrolysis ratio (GlcNAc6S: GalNAc6S)** | **Hydrolysis with BT3333$^{6S\text{-}GalNAc}$ (h$^{-1}$)** | **Hydrolysis with BT4656$^{6S\text{-}GlcNAc/GlcNS}$ (h$^{-1}$)** | **Enzymatic hydrolysis ratio(GlcNAc6S: GalNAc6S)** |
| **6S-GalNAc** | 0.36 | 0.9:1 | 2.5 | - | 121:1 |
| **6S-GlcNAc** | 0.33 | | - | 302 | |

**Table S9**

Primers utilised to generate genetic constructs in the pET28b vector.

| Primer name | Primer direction | Sequence |
|---|---|---|
| BT1918 | Foward | ggaatagctagccaaaaacaaccgcacattatctta |
| | Reverse | gagatactcgagttatcgcttttctttcggatagtt |
| BT1918$^{S73C}$ | Foward | cttcgtgtcccagctgcacccctgcccgtg |
| | Reverse | cacgggcaggggtgcagctgggacacgaag |
| BT1918$^{Y94A}$ | Foward | catggcttgttaggagccggtaaagttgcccccaagtac |
| | Reverse | gggggcaactttaccggctcctaacaagccatgatgcca |
| BT1918$^{Y94F}$ | Foward | catggcttgttaggatttggtaaagttgcccccaagtac |
| | Reverse | gggggcaactttaccaaatcctaacaagccatgatgcca |
| BT1918$^{R143A}$ | Foward | ctggatgaaagcggagcccgggaagatccgaatttcatc |
| | Reverse | attcggatcttcccgggctccgctttcatccagcagcgt |
| BT1918$^{W173A}$ | Foward | gaaaccggaatcggagccaacgatcacggagcagctacc |
| | Reverse | tgctccgtgatcgttggctccgattccggtttcatccgg |
| BT1918$^{N174A}$ | Foward | accggaatcggatgggccgatcacggagcagctacctat |
| | Reverse | agctgctccgtgatcggcccatccgattccggtttcatc |
| BT1918$^{R327A}$ | Foward | gatcactaccactgggccaagacttacccgtatgaagga |
| | Reverse | atacgggtaagtcttggcccagtggtagtgatctcccat |
| BT1918$^{Y408A}$ | Foward | gaacatgccacttgcgccagtgatgacaactattggtgt |
| | Reverse | atagttgtcatcactggcgcaagtggcatgttccaggtc |
| BT1918$^{Y408A}$ | Foward | gaacatgccacttgctttagtgatgacaactattggtgt |
| | Reverse | atagttgtcatcactaaagcaagtggcatgttccaggtc |
| BT3057 | Foward | ggaatagctagccaggaacgcccgaatattattgtct |
| | Reverse | gagatactcgagttaaaagtaaacaaaaggcttcaa |
| BT3057$^{S88C}$ | Foward | cgcacagagcgtgtgctccccttccagagc |
| | Reverse | gctctggaaggggagcacacgctctgtgcg |
| BT3057$^{W109A}$ | Foward | catggagtcactaacgccattaatgctgagagcaacaac |
| | Reverse | gctctcagcattaatggcgttagtgactccatgcctggt |
| BT3057$^{H182A}$ | Foward | ggttccggcattggagcccccggcagctattatggcgaa |
| | Reverse | ataatagctgccgggggctccaatgccggaacctgcgat |
| BT3057$^{H423A}$ | Foward | cacttccctcacgcagcccgcggcagctacttcaccacc |
| | Reverse | gaagtagctgccgcgggctgcgtgagggaagtgcatcag |
| BT3796 | Foward | ggaatagctagccaaacacagaagtctgcccagaca |
| | Reverse | gagatactcgagttacttaatttcgtccggccaagg |
| BT3796A$^{S88C}$ | Foward | gccagcagtatctgctcccccacccgatg |
| | Reverse | catcgggtgggggagcagatactgctggc |
| BT3796A$^{W109A}$ | Foward | caccgggtaaccaacgccacgctgcaaaagaacacaaag |
| | Reverse | gttcttttgcagcgtggcgttggttacccggtggcgtgc |
| BT3796A$^{W332A}$ | Foward | gccgccgaatcgtatgcccgtgacggcaaactgcacacc |
| | Reverse | cagtttgccgtcacgggcatacgattcggcggcaagtcc |
| BT3796A$^{W431A}$ | Foward | aatatgcccaacaacgccggaaacgacggtccgggaatc |
| | Reverse | cggaccgtcgtttccggcgttgttgggcatattccagaa |
| BT1624 | Foward | ggaatagctagcaaacaaacccagcgtccgaatgtt |
| | Reverse | gagatactcgagttatagtggcagaccgtagcgtcc |
| BT1624$^{S79C}$ | Foward | cagtgcagctgctacctgcacgccttcccgctatgcg |
| | Reverse | cgcatagcgggaaggcgtgcaggtagcagctgcactg |

| | | |
|---|---|---|
| BT1624$^{D170A}$ | Foward | atggcagctacgggcgcccgtgttccttgtgtatttgta |
| | Reverse | tacacaaggaacacgggcgcccgtagctgccatgataaa |
| BT1624$^{R171A}$ | Foward | gcagctacgggcgacgccgttccttgtgtatttgtagag |
| | Reverse | aaatacacaaggaacggcgtcgcccgtagctgccatgat |
| BT1624$^{H220A}$ | Foward | catccttctcacggagccgaccagagtatcgtgaacggt |
| | Reverse | cacgatactctggtcggctccgtgagaaggatgcatttt |
| BT1624$^{D336A}$ | Foward | ggtccggtagtggacgccggatataaggatcaggcagtg |
| | Reverse | ctgatccttatatccggcgtccactaccggaccgttatc |
| BT1624$^{Y360A}$ | Foward | tatcgcggaggaaaggccagttcgttcgaagcgggtaca |
| | Reverse | cgcttcgaacgaactggcctttcctccgcgataaatgcc |
| BT1624$^{Y360F}$ | Foward | tatcgcggaggaaagtttagttcgttcgaagcgggtaca |
| | Reverse | cgcttcgaacgaactaaactttcctccgcgataaatgcc |
| BT3333 | Foward | ggaatagctagcaatcagaaagtacaaaaagacaac |
| | Reverse | gagatactcgagttatcttgtctttaccgattcaag |
| BT3333$^{S83C}$ | Foward | gcaacttccgccacctgcacccccttcccgcttc |
| | Reverse | gaagcgggaagggggtgcaggtggcggaagttgc |
| BT3333$^{W464}$ | Foward | gccatcgaatactggacaaaaatggaactggc |
| | Reverse | ttccatttttgtccagtattcgatggcgggggcg |
| BT3333$^{\Delta Y463/W464}$ | Foward | ttcgatggcgggggcgtcactcgg |
| | Reverse | acaaaaatggaactgggcaatgac |
| BT3109 | Foward | ggaatagctagcgaagaaaagccgaatgtgattgtc |
| | Reverse | gagatactcgagttattttaatggttcttcttccac |
| BT3109$^{S73C}$ | Foward | gctacttcggctacttgcacccccagtcgttatgc |
| | Reverse | gcataacgactgggggtgcaagtagccgaagtagc |
| BT3109 $^{\Delta CT}$ | Foward | ctcactacgataaaatccatctgt |
| | Reverse | taactcgagcaccaccaccaccac |
| BT3177 | Foward | ggaatagctagccagaagccactgaacatcgtctat |
| | Reverse | gagatactcgagttattccttgtctctctcagggga |
| BT3177$^{S79C}$ | Foward | gtggctaattctctgtgtggacccagccgtgcc |
| | Reverse | ggcacggctgggtccacacagagaattagccac |
| BT3177$^{Q147A}$ | Foward | gaaatagttcccggtgccggcgattactacaatccggat |
| | Reverse | attgtagtaatcgccggcaccgggaactatttcccagta |
| BT3177$^{R202A}$ | Foward | cacaaagcgattcatgccaactggttggcagatacgtgt |
| | Reverse | atctgccaaccagttggcatgaatcgctttgtgatgaat |
| BT3177$^{Y250A}$ | Foward | gatatggatatgatcgccgatcttaaaatgctgcgtcct |
| | Reverse | cagcattttaagatcggcgatcatatccatatctttggc |
| BT3177$^{Y250F}$ | Foward | gatatggatatgatctttgatcttaaaatgctgcgtcct |
| | Reverse | cagcattttaagatcaaagatcatatccatatctttggc |
| BT3177$^{D361A}$ | Foward | gagcatggttggtttgccaaacgtttcatgtacgaagag |
| | Reverse | gtacatgaaacgtttggcaaaccaaccatgctcacccat |
| BT3177$^{R363A}$ | Foward | ggttggtttgacaaagccttcatgtacgaagagtccatg |
| | Reverse | ctcttcgtacatgaaggctttgtcaaaccaaccatgctc |
| BT3177$^{E444A}$ | Foward | tatgaatatcctgccgcccacatggtgaaacgccattac |
| | Reverse | gcgtttcaccatgtgggcggcaggatattcatagaaatg |
| BT3177$^{H445A}$ | Foward | gaatatcctgccgaagccatggtgaaacgccattacggc |
| | Reverse | atggcgtttcaccatggcttcggcaggatattcatagaa |

**Table S10**

Melting temperatures of wildtype and mutant proteins.

| Protein | $T_m$ |
|---|---|
| $BT1918^{S73C}$ | 42.8 ± 0.21 |
| $BT1918^{Y94A}$ | 44.9 ± 0.51 |
| $BT1918^{Y94F}$ | 43.2 ± 0.19 |
| $BT1918^{R143A}$ | 41.1 ± 0.11 |
| $BT1918^{W173A}$ | 41.1 ± 0.25 |
| $BT1918^{N174A}$ | 40.0 ± 0.18 |
| $BT1918^{R327A}$ | 40.8 ± 0.40 |
| $BT1918^{Y408A}$ | 40.9 ± 0.20 |
| $BT1918^{Y408A}$ | 45.3 ± 0.44 |
| $BT3057^{S88C}$ | 60.3 ± 0.80 |
| $BT3057^{W109A}$ | 62.7 ± 0.40 |
| $BT3057^{H182A}$ | 62.11 ± 0.64 |
| $BT3057^{H423A}$ | 58.6 ± 0.63 |
| $BT3796A^{S88C}$ (1st melt curve) | 42.0 ± 0.12 |
| $BT3796A^{W109A}$ (1st melt curve) | 60.0 ± 0.07 |
| $BT3796A^{W332A}$ (1st melt curve) | 56.2 ± 0.35 |
| $BT3796A^{W431A}$ (1st melt curve) | 44.6 ± 0.14 |
| $BT3796A^{S88C}$ (2nd melt curve) | 55.0 ± 0.01 |
| $BT3796A^{W109A}$ (2nd melt curve) | NT* |
| $BT3796A^{W332A}$ (2nd melt curve) | NT* |
| $BT3796A^{W431A}$ (2nd melt curve) | 57.2 ± 0.03 |
| $BT1624^{S79C}$ | 56.3 ± 0.08 |
| $BT1624^{D170A}$ | 48.2 ± 0.05 |
| $BT1624^{R171A}$ | 59.7 ± 0.04 |
| $BT1624^{H220A}$ | 46.8 ± 0.06 |
| $BT1624^{D336A}$ | 49.0 ± 0.08 |
| $BT1624^{Y360A}$ | 48.7 ± 0.03 |
| $BT1624^{Y360F}$ | 50.3 ± 0.11 |
| $BT3333^{S83C}$ | 57.7 ± 0.07 |
| $BT3333^{W464}$ | 57.7 ± 0.07 |
| $BT3333^{\Delta Y463/W464}$ | 54.6 ± 0.09 |
| $BT3109^{S73C}$ | 53.4 ± 0.02 |
| $BT3109^{\Delta CT}$ | 52.8 ± 0.06 |

All assays were conducted in technical triplicates and were carried out in 100 mM BTP pH 7.0 with 150 mM NaCl using 5 μM of the appropriate protein. NT indicates not tested as the melting curves could no longer be separated and analysed separately.

**Table S11**

Melting temperatures of wildtype and mutant proteins in the presence and absence of product.

| **Protein** | **Melting temperature ($T_m$)** | | | | | | |
|---|---|---|---|---|---|---|---|
| | No ligand | **1 mM Gal** | **10 mM Gal** | **100 mM Gal** | **1 mM GalNAc** | **10 mM GalNAc** | **100 mM GalNAc** |
| BT3057$^{S88C}$ | 60.3 ± 0.80 | 58.3 ± 0.09 | 58.2 ± 0.15 | 59.3 ± 0.07 | 58.1 ± 0.11 | 58.2 ± 0.07 | 58.9 ± 0.12 |
| BT3057$^{W109A}$ | 62.7 ± 0.40 | NT | NT | NT | NT | NT | NT |
| BT3057$^{H182A}$ | 62.11 ± 0.64 | NT | NT | NT | NT | NT | NT |
| BT3057$^{H423A}$ | 58.6 ± 0.63 | NT | NT | NT | NT | NT | NT |
| BT3796A$^{S88C}$ (1$^{st}$ melt curve) | 42.0 ± 0.12 | 42.1 ± 0.05 | 42.3 ± 0.05 | 43.8 ± 0.21 | 42.1 ± 0.01 | 42.0 ± 0.12 | 42.3 ± 0.37 |
| BT3796A$^{S88C}$ (2$^{nd}$ melt curve) | 55.0 ± 0.01 | 55.0 ± 0.02 | 55.0 ± 0.02 | 55.1 ± 0.02 | 54.9 ± 0.01 | 54.9 ± 0.03 | 54.9 ± 0.05 |
| BT1624$^{S79C}$ | 56.3 ± 0.08 | NT | NT | 60.6 ± 0.30 | NT | NT | NT |
| BT1624$^{D170A}$ | 48.2 ± 0.05 | NT | NT | 49.6 ± 0.02 | NT | NT | NT |
| BT1624$^{R171A}$ | 59.7 ± 0.04 | NT | NT | 60.8 ± 0.05 | NT | NT | NT |
| BT1624$^{H220A}$ | 46.8 ± 0.06 | NT | NT | 48.4 ± 0.04 | NT | NT | NT |
| BT1624$^{D336A}$ | 49.0 ± 0.08 | NT | NT | 50.2 ± 0.04 | NT | NT | NT |
| BT1624$^{Y360A}$ | 48.7 ± 0.03 | NT | NT | 50.6 ± 0.04 | NT | NT | NT |
| BT1624$^{Y360F}$ | 50.3 ± 0.11 | NT | NT | 52.3 ± 0.04 | NT | NT | NT |
| BT3333$^{S83C}$ | 57.7 ± 0.07 | 57.7 ± 0.02 | 57.9 ± 0.11 | 59.0 ± 0.08 | 58.8 ± 0.13 | 61.2 ± 0.18 | 64.5 ± 0.26 |
| BT3333$^{W464}$ | 57.7 ± 0.07 | 57.8 ± 0.03 | 58.0 ± 0.04 | 59.2 ± 0.06 | 57.7 ± 0.11 | 58.7 ± 0.05 | 61.2 ± 0.03 |
| BT3333$^{\Delta Y463/W464}$ | 54.6 ± 0.09 | 53.6 ± 0.02 | 53.9 ± 0.12 | 55.4 ± 0.21 | 53.7 ± 0.05 | 54.2 ± 0.03 | 56.7 ± 0.20 |
| BT3109$^{S73C}$ | 53.4 ± 0.02 | 54.0 ± 0.09 | 55.0 ± 0.20 | 56.1 ± 0.11 | 53.2 ± 0.04 | 53.4 ± 0.08 | 53.7 ± 0.01 |
| BT3109 $^{\Delta CT}$ | 52.8 ± 0.06 | 53.2 ± 0.25 | 53.1 ± 0.07 | 53.8 ± 0.13 | 52.7 ± 0.04 | 52.9 ± 0.05 | 53.6 ± 0.3 |

All assays were conducted in technical triplicates and were carried out in 100 mM BTP pH 7.0 with 150 mM NaCl using 5 μM of the appropriate protein.

**Table S12**

Data collection and refinement statistic for crystallographic data of S1_46 and S1_16 sulfatases.

| | BT1918$^{3S\text{-}Gal}$ | BT3057$^{4S\text{-}Gal/GalNAc}$ | BT3796$^{4S\text{-}Gal/GalNAc}$ |
|---|---|---|---|
| Beamline | I04 | I04-1 | I04-1 |
| Date | 07/08/18 | 28/07/18 | 28/07/18 |
| Wavelength (Å) | 0.98 | 0.92 | 0.92 |
| Resolution (Å) | 48.1-1.95 (2.0-1.95) | 65.43-1.91 (1.94-1.91) | 81.48-1.50 (1.53-1.50) |
| Space group | $P3_22_1$ | $P2_1$ | $P2_1$ |
| Unit-cell parameters | | | |
| a, b, c (Å) | 56.9, 56.9, 279.8 | 66.2, 90.8, 94.2 | 73.2, 88.0, 87.9 |
| α , β γ, (°) | 90, 90, 120 | 90, 98.5, 90 | 90, 112.1, 90 |
| No. of measured reflections | 404335 (27844) | 164475 (8211) | 314826 (15779) |
| No. of independent reflections | 39105 (2675) | 85409 (4269) | 166081 (8213) |
| Completeness (%) | 100 (100) | 99.6 (94.6) | 99.7 (99.8) |
| Redundancy | 10.3 (10.4) | 1.9 (1.9) | 1.9 (1.9) |
| <I>/<σ(I)> | 5.8 (1.5) | 8.8 (2.2) | 7.0 (1.5) |
| CC(1/2) | 0.990 (0.803) | 0.992 (0.39) | 0.996 (0.577) |
| Refinement statistics* | | | |
| $R_{work}/R_{free}$ (%) | 19/22 | 18/22 | 14/18 |
| No. of non-H atoms | | | |
| No. of protein, atoms | 3801 | 7584 | 7776 |
| No. of solvent atoms | 166 | 405 | 755 |
| No. of ligand atoms | 23 | 76 | 96 |
| R.m.s. deviation from ideal values | | | |
| Bond angle (°) | 1.52 | 1.6 | 1.5 |
| Bond length (Å) | 0.008 | 0.008 | 0.008 |
| Average B factor (Å$^2$) | | | |
| Protein | 31.8 | 25.6 | 17.8 |
| Solvent | 30.7 | 27.8 | 29.7 |
| Ligand | 25.7 | 19.7 | 33.5 |
| Ramachandran plot$^+$, most favoured regions (%) | 96.4 | 96.5 | 97.5 |
| PDB code | 7OZ8 | 7OZ9 | 7OZA |

Values in parenthesis are for the highest resolution shell. $R_{free}$ was calculated using a set (5%) of randomly selected reflections that were excluded from refinement.

**Table S13**

Data collection and refinement statistics for crystallographic data of S1_15 and S1_11 sulfatases.

| | BT1624$^{6S-Gal/GalNAc}$ | BT3109$^{6S-Gal/GalNAc}$ | BT4631$^{6S-Gal/GalNAc}$ | BT3177$^{6S-GlcNAc}$ |
|---|---|---|---|---|
| Beamline | I04 | I03 | PROXIMA-SOLEIL | I04-1 |
| Date | 07/08/18 | 14/12/19 | 12/09/20 | 28/07/18 |
| Wavelength (Å) | 0.98 | 0.98 | 0.98 | 0.92 |
| Resolution (Å) | 50.5-2.7 (2.78-2.70) | 47.09-1.75 (1.78-1.75) | 109.57-1.35 (1.37-1.35) | 35.83-0.97 (0.99-0.97) |
| Space group | $P2_1$ | $P2_1$ | $P22_12_1$ | $P2_1$ |
| Unit-cell parameters | | | | |
| a, b, c (Å) | 86.4, 124.7, 97.1 | 52.2, 91.8, 52.7 | 49.92, 109.57, 117.53 | 42.1, 69.8, 96.3 |
| α , β γ, (°) | 90, 95.3, 90 | 90, 116.6, 90 | 90,90,90 | 90, 97.6, 90 |
| No. of measured reflections | 207968 (16852) | 153801 (8518) | 1739450 (46455) | 553797 (22206) |
| No. of independent reflections | 56316 (4596) | 44392 (2410) | 140145 (6050) | 298423 (12666) |
| Completeness (%) | 100 (99.9) | 99.5 (99) | 98.8 (88.1) | 91.9 (79.0) |
| Redundancy | 3.7 (3.7) | 3.5 (3.5) | 12.4 (7.7) | 1.9 (1.8) |
| $\langle I\rangle/\langle\sigma(I)\rangle$ | 3.0 (1.4) | 8.8 (1.2) | 25.4 (4.5) | 8.8 (1.5) |
| CC(1/2) | 0.927 (0.662) | 0.997 (0.493) | 0.999 (0.955) | 0.996 (0.596) |
| Refinement statistics* | | | | |
| $R_{work}/R_{free}$ (%) | 23/26 | 18/22 | 12/14 | 12/14 |
| No. of non-H atoms | | | | |
| No. of protein, atoms | 14902 | 3873 | 4074 | 4476 |
| No. of solvent atoms | 267 | 186 | 599 | 595 |
| No. of ligand atoms | 64 | 16 | 28 | 38 |
| R.m.s. deviation from ideal values | | | | |
| Bond angle (°) | 1.46 | 1.49 | 1.48 | 1.50 |
| Bond length (Å) | 0.007 | 0.008 | 0.008 | 0.008 |
| Average B factor (Å$^2$) | | | | |
| Protein | 17.9 | 29.2 | 13.8 | 11.3 |
| Solvent | 7.3 | 31.4 | 28.9 | 24.2 |
| Ligand | 18.4 | 21.0 | 41.8 | 7.53 |
| Ramachandran plot$^+$, most favoured regions (%) | 96.6 | 96.5 | 95.4 | 96.3 |
| PDB code | 7OZE | 7OZC | 7P26 | 7P24 |

Values in parenthesis are for the highest resolution shell. $R_{free}$ was calculated using a set (5%) of randomly selected reflections that were excluded from refinement.

1 Luis, A. S. *et al.* A single bacterial sulfatase is required for metabolism of colonic mucin <em>O</em>-glycans and intestinal colonization by a symbiotic human gut bacterium. *bioRxiv*, 2020.2011.2020.392076, doi:10.1101/2020.11.20.392076 (2020).

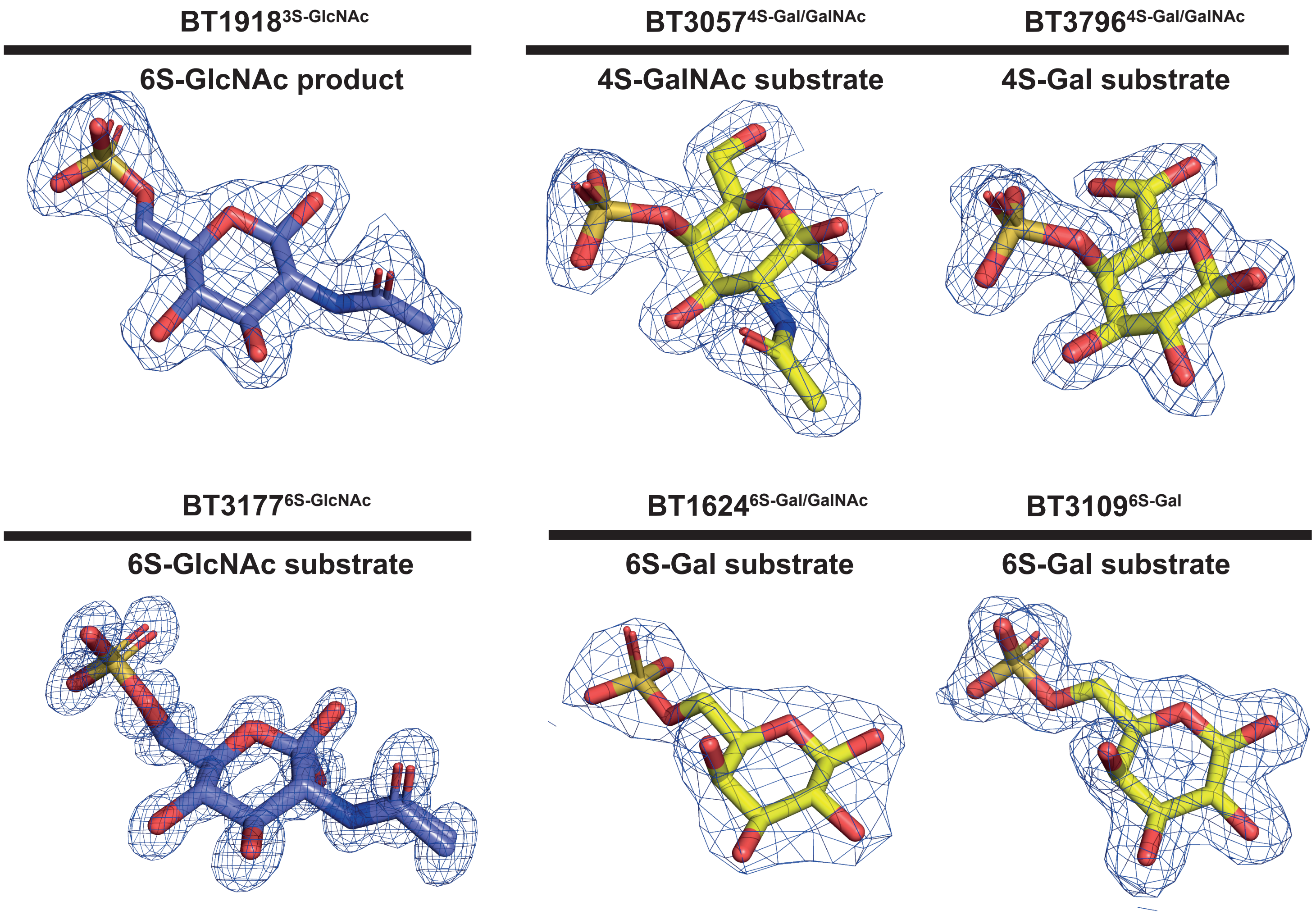


**Supplemental Figure 1. Extracted ligands and their electron density maps.**
The 2mFobs-Fc maps are shown contoured at 1σfor all substrates and products co-crystallised with their respective sulfatase.

Supplementary Figure 1

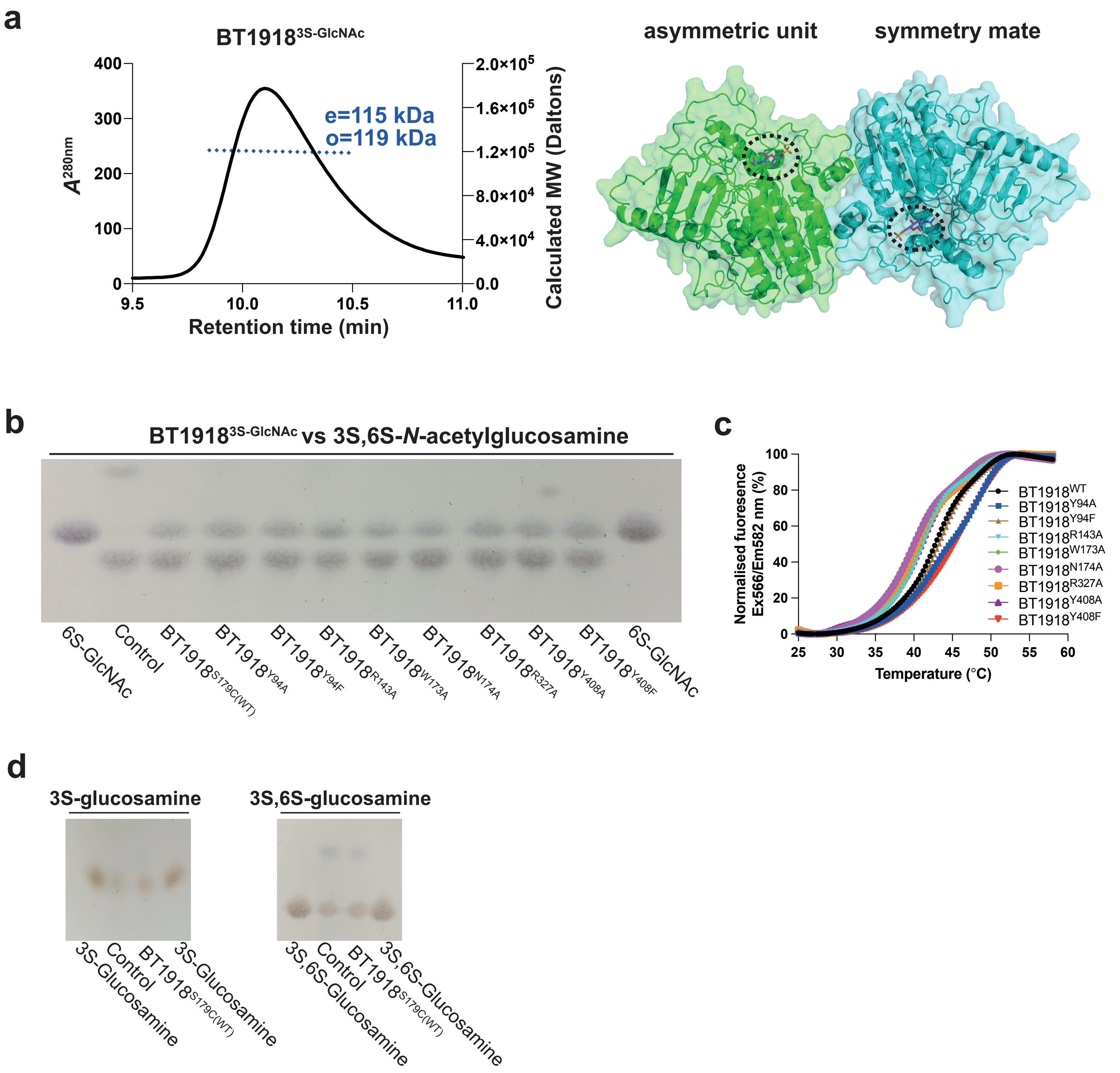


**Supplemental Figure 2. Biophysical and kinetics analysis of BT19183S-GlcNAc**

**a,** Right panel shows chromatogram of size exclusion chromatography coupled to light scattering for BT1918$^{3S\text{-}GlcNAc}$ demonstrating the presence of a dimer(e= expected molecular mass; o=observed molecular mass). Left panel shows surface representation of the dimer of BT1918$^{3S\text{-}GlcNAc}$; the active site is highlighted with a black dashed circle. **b,** Thin layer chromatography analysis of BT1918$^{3S\text{-}GlcNAc}$ and its mutants versus its 3S, 6S-N-acetylglucosamine substrate. Assays were ran for 48 h, at 37ºC, deploying 6 mM substrate and 5 µM enzyme. All assays contained 3 mM HEPES pH 7.0, 45 mM NaCl, and 5 mM $CaCl_2$. **c,** DSF analysis showing the stability of the mutant proteins of BT1918$^{3S\text{-}GlcNAc}$ with respect to the wildtype (WT). **d,** Thin layer chromatography analysis of BT1918$^{3S\text{-}GlcNAc}$ versus 3S-glucosamine and 3S,6S-glucosamine. Assays were ran for 48 h, at 37ºC, deploying 6 mM substrate and 5 µM enzyme. All assays contained 3 mM HEPES pH 7.0, 45 mM NaCl, and 5 mM $CaCl_2$.

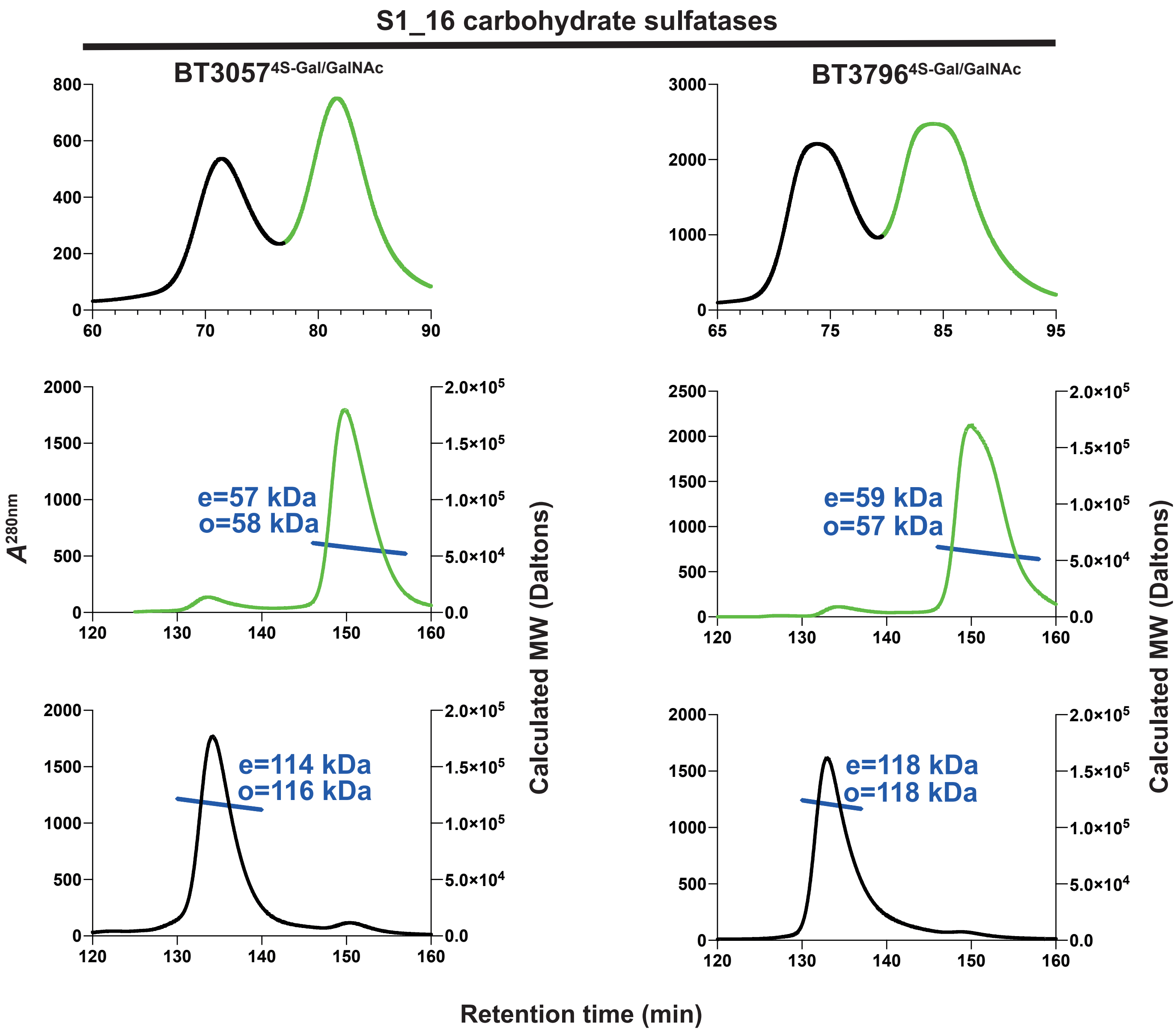


**Supplemental Figure 3. Gel chromatography and light scattering of S1_16 sulfatases.**
Top panels are size exclusion chromatograms whilst, the middle and bottom panels are size exclusion coupled to light scattering chromatograms for the monomer (green) and dimer (black) peaks purified from size exclusion. e= expected molecular mass; o=observed molecular mass.

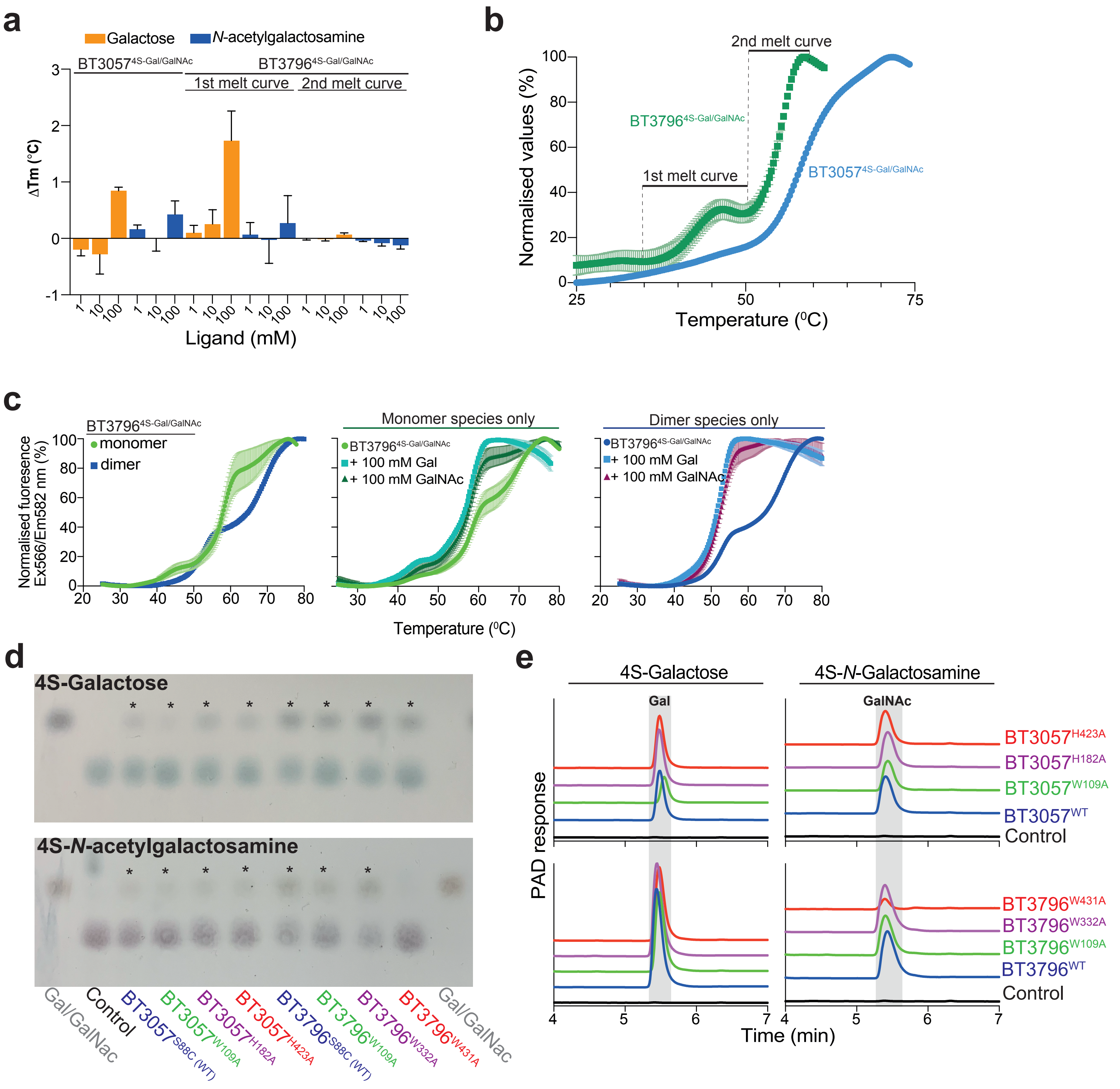


**Supplemental Figure 4. Activity and stability analysis of S1_16 sulfatases and their mutant variants.**
**a,** DSF analysis of the effects of galactose and N-acetyl galactosamine on thermostability with a positive shift indicative of binding. **b,** Normalised DSF melt curves of $BT3057^{4S\text{-}Gal/GalNAc}$ and $BT3796^{4S\text{-}Gal/GalNAc}$. **c,** DSF melt curves of the purified monomer and dimer species (left), monomer species in the presence of galactose and N-acetyl galactosamine (middle), and the dimer species in the presence of galactose and *N*-acetyl galactosamine (right). **d,** Thin layer chromatography (TLC) analysis of wildtype (WT) and mutant S1_16 sulfatases. Asterisks are placed above lanes where activity is observed. e, High pressure anion exchange chromatography (HPAEC) of wildtype (WT) and mutant S1_16 sulfatases. A grey block highlights the produced product. Both TLC and HPAEC reactions utilised 6 mM substrate and 1 mM enzyme, except for W109A variants where 10 µM was deployed, with 3 mM HEPES, 45 mM NaCl, and 5 mM CaCl2 over a 48 h period at 37ºC.

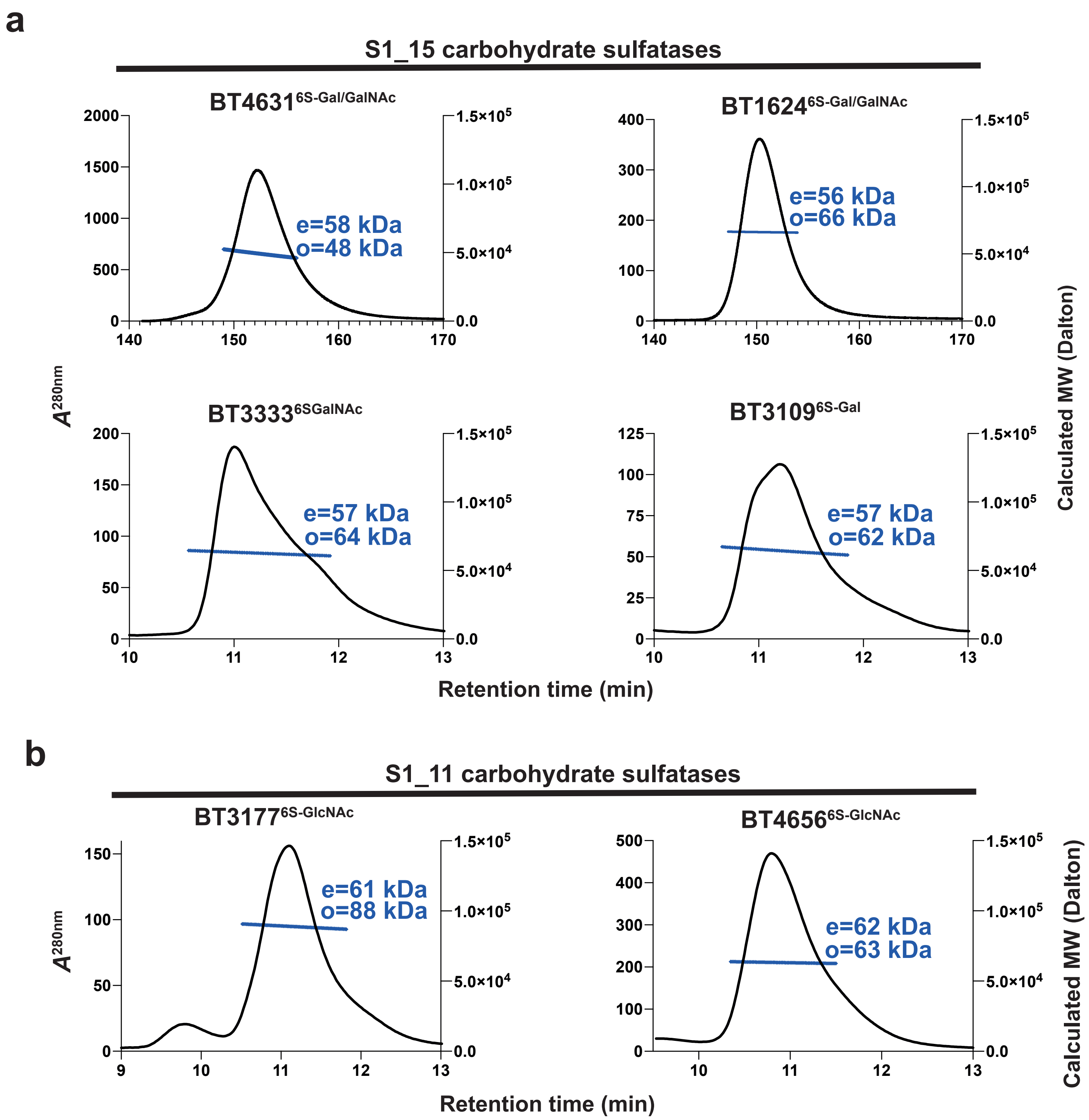


**Supplemental Figure 5. Gel chromatography and light scattering of S1_15 and S1_11 sulfatases.** a, Chromatograms of size exclusion chromatography coupled to light scattering for S1_15 subfamily members; b, Chromatograms of size exclusion chromatography coupled to light scattering for S1_11 subfamily members. e= expected molecular mass; o=observed molecular mass.

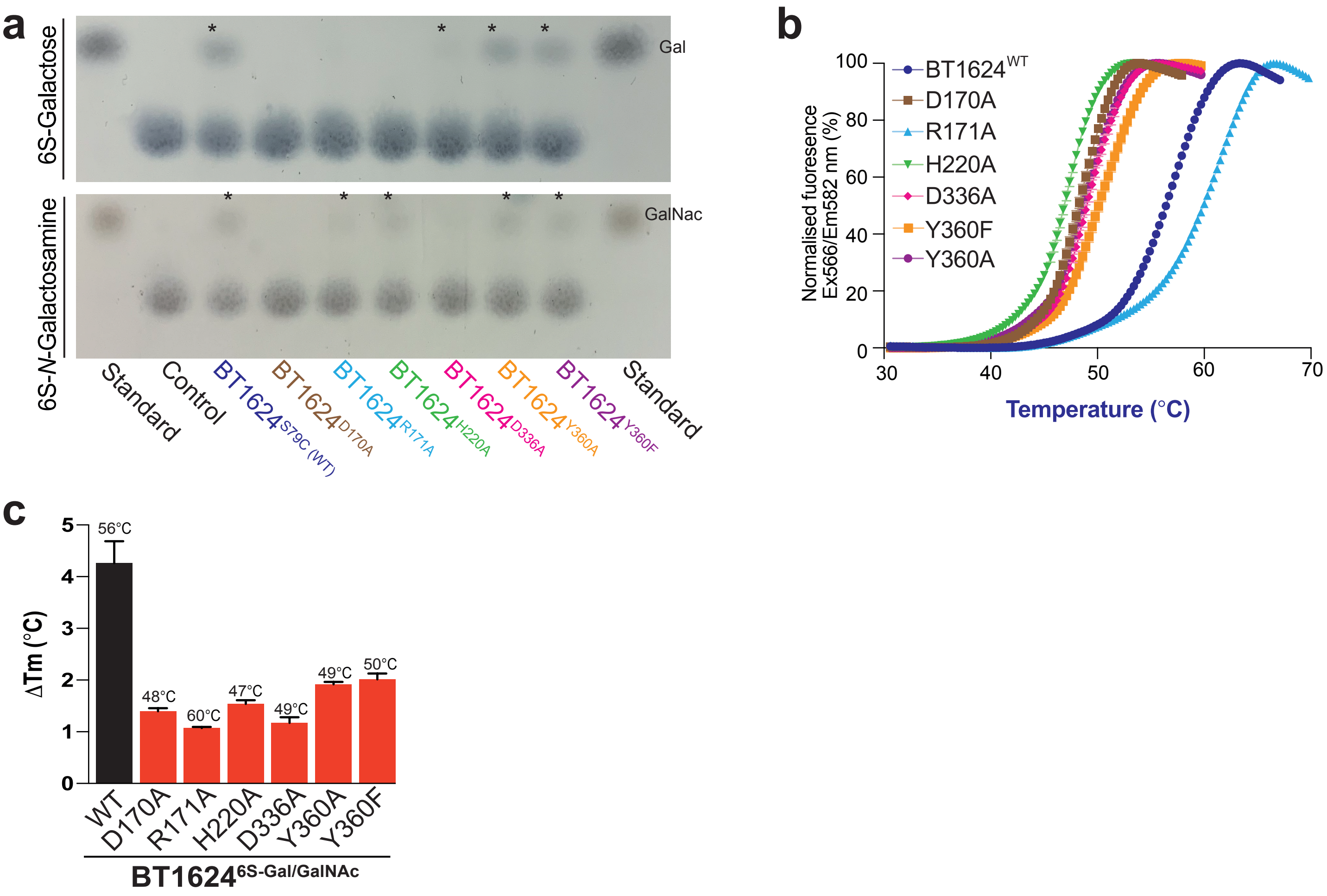


**Supplemental Figure 6. Analysis of the activity and stability of BT1624$^{6S\text{-}Gal/GalNAc}$ and its mutant variants.**

**a,** Thin layer chromatography (TLC) analysis of wildtype (WT) BT1624$^{6S\ Gal/GalNAc}$ and its mutants. Asterisks are placed above lanes where activity is observed. Both TLC reactions utilised 6 mM substrate and 5 µM enzyme, with 3 mM HEPES, 45 mM NaCl, and 5 mM $CaCl_2$ over a 48 h period at 37ºC. **b,** DSF analysis showing the stability of the mutant proteins of BT1624$^{6S\text{-}Gal/GalNAc}$ with respect to the wildtype. **c,** DSF analysis of the effects of alanine scanning on the ability of BT1624$^{6S\text{-}Gal/GalNAc}$ to bind galactose with the Tm of the protein shown atop the bar. The experiments were performed with 5 µM of protein and 324 mM of galactose in 100 mM BTP and 150 mM NaCl.

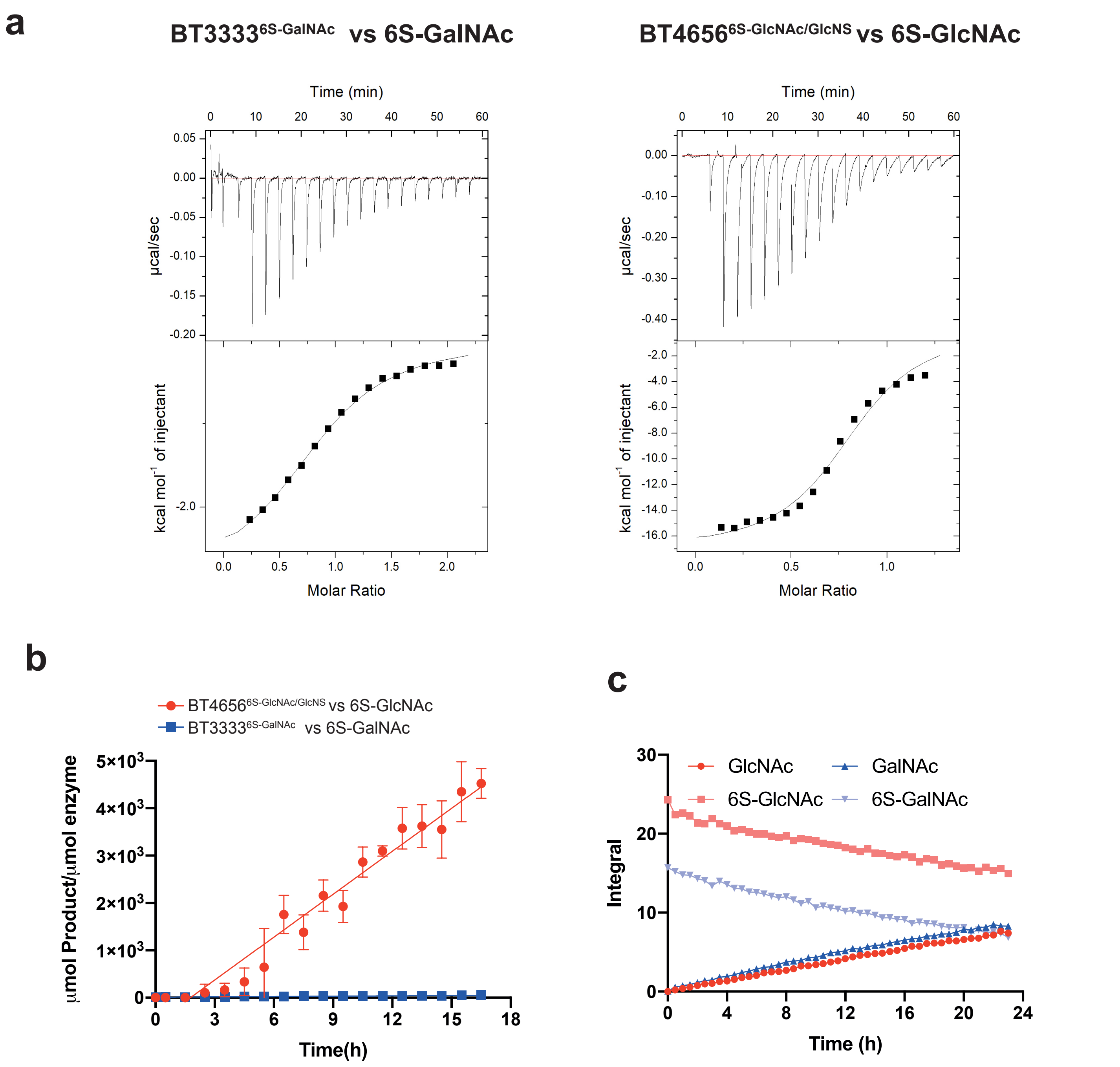


**Supplemental Figure 7. Biochemical comparison of S1_15 BT3333$^{\text{6S-GalNAc}}$ and S1_11 sulfatase BT4656$^{\text{6S-GlcNAc/GlcNS}}$**

**a,** Isothermal titration calorimetry traces of inactive serine variants of BT33336S-GalNAc and BT4656$^{\text{6S-GlcNAc/GlcNS}}$ binding to their respective 6S-GalNAc and 6S-GlcNAc substrates. **b,** Specific activity of BT3333$^{\text{6S-GalNAc}}$ and BT4656$^{\text{6S-GlcNAc}}$ against 0.5 mM their respective 6S-GalNAc and 6S-GlcNAc substrates, monitored by 1D H-NMR. c, Acid hydrolysis of 10 mM 6S-GalNAc and 6S-GlcNAc in the presence of 250 mM HCl at 90°C.

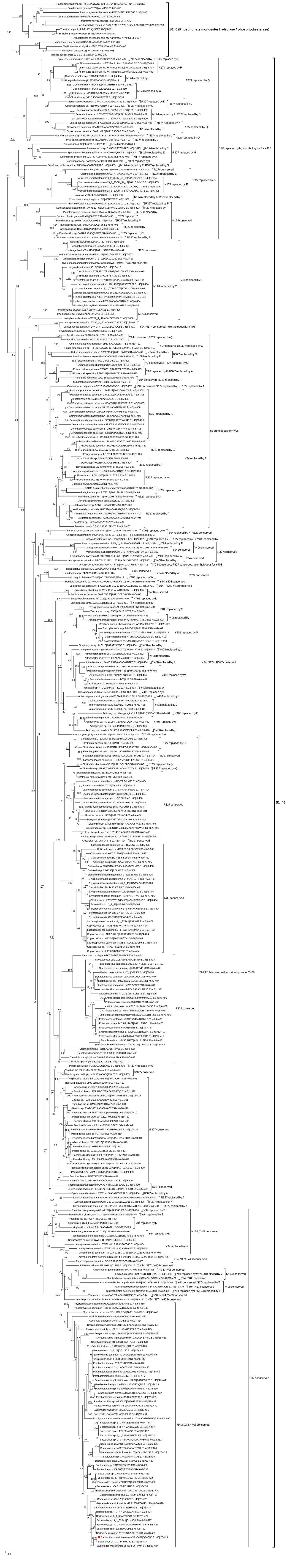
S1_0 (Phosphonate monoester hydrolase / phosphodiesterase)
S1_46

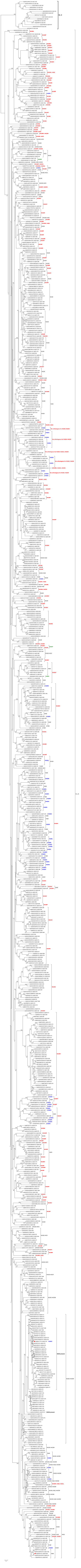

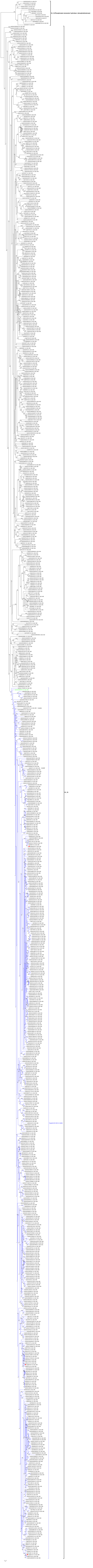
S1_0 (Phosphonate monoester hydrolase / phosphodiesterase)
S1_15
Triad D170 / R171 / H220
H220F
0.2

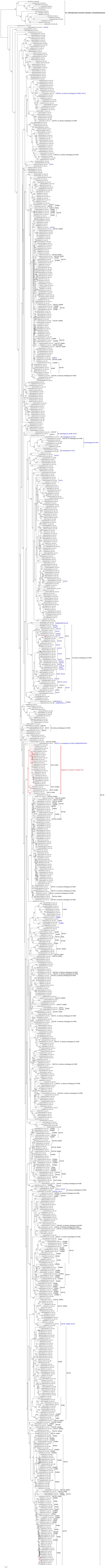
